# A novel mechanosensitive channel controls osmoregulation, differentiation and infectivity in *Trypanosoma cruzi*

**DOI:** 10.1101/498469

**Authors:** N Dave, U Cetiner, D Arroyo, J Fonbuena, M Tiwari, P Barrera, N Lander, A Anishkin, S Sukharev, V Jimenez

## Abstract

*Trypanosoma cruzi*, the causative agent of Chagas disease, undergoes drastic morphological and biochemical modifications as it passes between hosts and transitions from extracellular to intracellular stages. The osmotic and mechanical aspects of these cellular transformations are not understood. Here we identify and characterize a novel mechanosensitive channel in T. *cruzi* (TcMscS) belonging to the superfamily of small conductance mechanosensitive channels (MscS). TcMscS is activated by membrane tension and forms a large pore permeable to anions, cations, and small osmolytes. The channel changes its location from the contractile vacuole complex in epimastigotes to the plasma membrane as the parasites develop into intracellular amastigotes. TcMscS knockout parasites show significant fitness defects, including increased cell volume, calcium dysregulation, impaired differentiation, and a dramatic decrease in infectivity. Our work provides mechanistic insights into components supporting pathogen adaptation inside the host thus opening the exploration of mechanosensation as a prerequisite of protozoan infectivity.

## Introduction

Mechanosensation is a universal characteristic of all cells, from bacteria to mammals [1, 2]. Mechanosensitive channels (MSCs) act as sensors (transducers) and often as effectors of mechano-responses [3] and their activation leads to the movement of ions and small osmolytes, which mediates regulatory volume responses and/or triggers downstream signaling cascades [4–8]. Thus, mechanosensation allows cells to respond to stress conditions, maintaining relatively constant volume and macromolecular excluded volume of the cytoplasm. In eukaryotic organisms mechanosensation is involved in shear stress sensing and stem cell differentiation [9–11], regulating the fate of mesenchymatic tissues such as myoblasts and osteoblasts [12, 13]. Importantly, mechanical cues regulate the growth of normal and tumoral tissue [14–16], allowing for the invasion of the basal lamina and increasing the metastasis of tumors [17, 18]. MSCs in prokaryotes have been postulated to participate in quorum sensing, biofilm formation and regulation of virulence factors [19–22]. Protozoan pathogens, like trypanosomes and apicomplexans, are subjected to mechanical forces associated with shear stress in the bloodstream [23, 24], extravasation [25, 26] and invasion of tissues required for establishing persistent infection [27–31]. While it is clear that mechanical cues modulate the parasite life cycle and behavior, the molecular mechanism underlying sensing and triggering of the responses required for survival is unknown.

*Trypanosoma cruzi*, a flagellated protozoan, is the causative agent of Chagas disease, an endemic pathology in Latin America where millions are infected [32]. Recent studies indicate global changes in the epidemiology of the disease resulting in an increased incidence in non-endemic regions of the world [33, 34]. Similar to other protozoan pathogens [35–37], *T. cruzi* completes its life cycle by alternating between mammalian hosts and insect vectors. The parasite undergoes transitions between intracellular and extracellular forms, with periods of exposure to the environment. Drastic morphological and biochemical changes are required to ensure its survival in different environments [35]. *T. cruzi* epimastigotes replicate in the intestinal lumen of the insect vector where they endure osmotic concentrations of rectal material up to 1000 mOsm/Kg [38]. The infective metacyclic trypomastigotes are released into the environment with the feces of the vector and are able to penetrate the skin layers of a mammalian host, disseminating via blood circulation. The highly motile flagellated trypomastigotes survive shear stress in the bloodstream and then penetrate into host cells where, in the relatively constant cytoplasmic environment, they transform into the round non-motile intracellular amastigotes. The differentiation back to the extracellular infective trypomastigotes requires substantial volume and shape changes, cytoskeletal rearrangement, and elongation of the flagella. Additionally, *T. cruzi* has a strong tropism toward muscle cells, establishing chronic infection in cardiomyocytes and smooth muscle of the gastrointestinal track [39]. Parasite survival and growth in contractile tissues, as well as developmental transitions from intracellular to extracellular forms, pose significant mechanical challenges for the parasites [40].

Multiple signaling pathways are activated during osmotic responses, invasion and differentiation [41–44] including cAMP and calcium-dependent cascades [45, 46], but the primary molecular sensors triggering these processes have yet to be identified. *T. cruzi* regulates its volume via the Contractile Vacuole Complex (CVC) [47, 48] a specialized organelle also present in other protozoans like *Paramecium, Dictyostelium and Chamydomonas*, which experience substantial osmotic gradients across their plasma membranes. The CVC actively collects excessive fluid from the cytoplasm and ejects it through exocytic mechanisms [49]. In *Chlamydomonas*, yeast, and various higher plants, the mechanisms of volume adjustment, stress relief, biogenesis, and maintenance of intracellular organelles such as plastids are regulated by mechanosensitive channels [50–53]. These channels belonging to the prokaryotic MscS family, act as ubiquitous turgor regulators and osmolyte release valves in bacteria [4, 54, 55] and Archaea [56]. Currently, there is no information about such tension-stabilizing components in protozoan CVCs.

Here, we report the identification and functional characterization of a novel mechanosensitive channel in *T. cruzi* (TcMscS), which is a member of a MscS-related branch of channels present in Trypanosomatids [57]. The sequence analysis predicts a two-transmembrane domain architecture with a unique C-terminal domain. During the parasite life-cycle the channel is developmentally targeted to CVC in the extracelullar motile stages and to the plasma membrane in the intracellular amastigote stage. When expressed in bacterial spheroplasts the channel gates directly by membrane tension and shows a slight selectivity for anions. Gene targeting by CRISPR/Cas methods generated knockout and knockdown parasites that exhibit impaired growth and inability to robustly regulate cell volume. These parasites also show abnormal calcium regulation and a dramatic decrease in differentiation rate and infectivity, supporting the essential role of mechanosensitive channels in *T. cruzi*. Our results provide evidence that in trypanosomes, TcMscS and its homologs are involved in the sensing of mechanical forces generated during tissue migration and cell penetration, stabilizing pressure gradients and relieving mechanical membrane stresses during cell volume regulation and developmental transitions.

## Results

### Sequence analysis and structural features of TcMscS predicted by homology model

Analysis of Trypanosoma databases (tritrypdb.org) revealed the presence of a putative MscS-like ion channel (TcCLB.504171.40) in the Esmeraldo-like haplotype of the *T. cruzi* CL Brener strain genome (TcMscS). This channel shares 94.5% identity at the protein level with the non Esmeraldo-like allele (TcCLB.509795.40) and has homologues in *T. brucei* (Tb427.10.9030) and *Leishmania major* (LmjF.36.5770), sharing 66% and 58% identity, respectively (Fig. 1A). Overall sequence conservation between TcMscS and *E. coli* MscS is low (25.4%), but the percentage of identity increases to 38% in the core comprising transmembrane helices TM2 and TM3. The full alignment of four protozoan and two bacterial sequences is shown in supplemental Figure S1. The ORF of TcMscS is 498 nucleotides, predicting a protein of 165 amino acids (17.8 kDa). Assuming that TcMscS follows the same oligomerization pattern as *E. coli* MscS [58], it is expected to form a heptamer. The refined consensus homology model based on several structure prediction servers (Fig. 1B) indicates the presence of two transmembrane domains, with a preceding N-terminal segment (red). The chain continues as TM2 (gold), crosses the membrane, makes a short loop and returns back as pore-forming TM3 (green). The gate of TcMscS is predicted to be formed by the ring of F64, which is in the precise homologous position with the gate-forming L109 of EcMscS.

**Figure 1.**
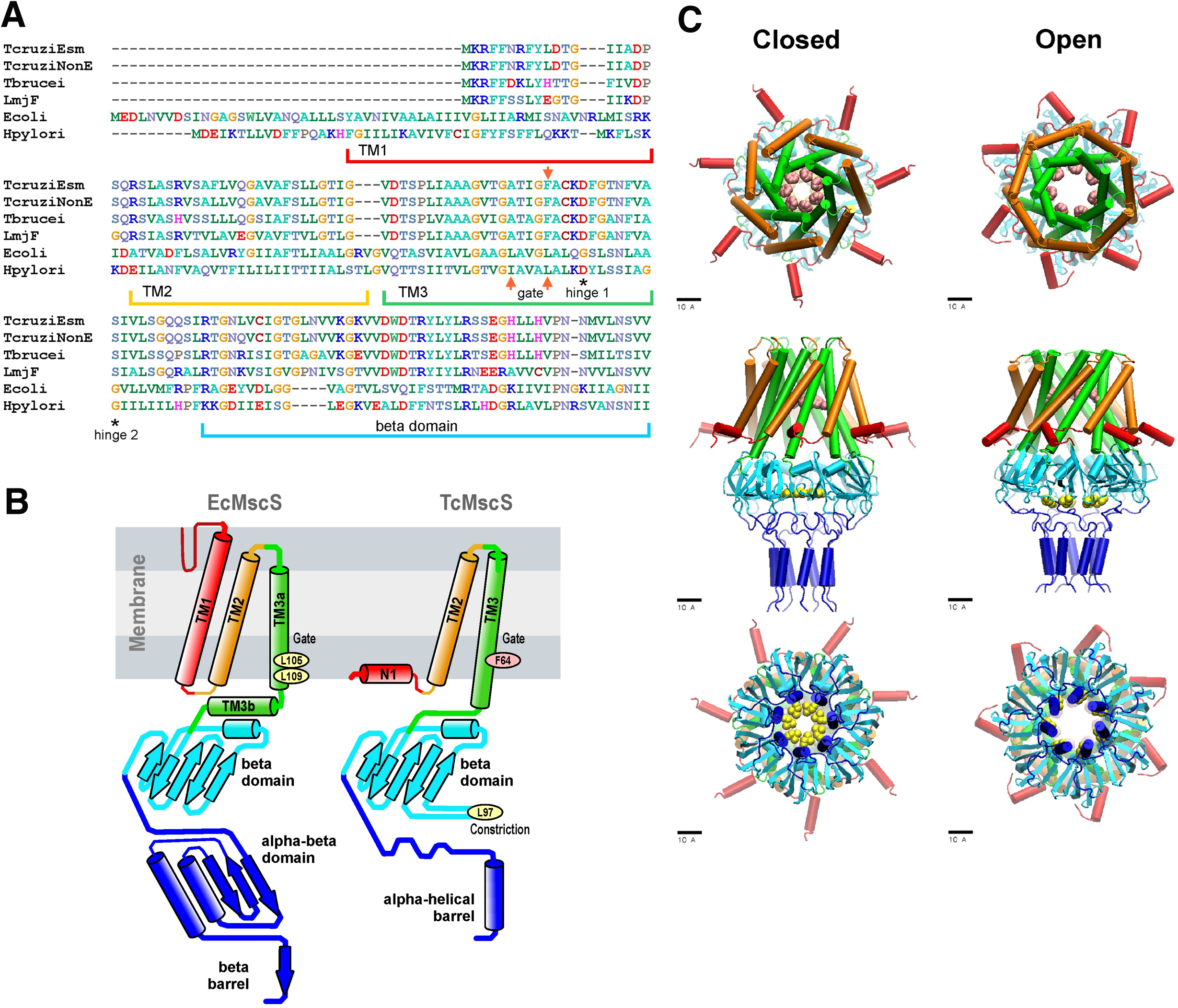
Sequence alignment and predictions of TcMscS structure by homology to EcMscS. **A.** Partial protein sequence alignment of four MscS-type channels from *T. cruzi T. cruzi* Esmeraldo-like (TcCLB.504171.40) and non-Esmeraldo-like (TcCLB.509795.40) haplotypes of the CL Brener reference strain, *T. brucei* (Tb427.10.9030) and *L. major* (LmjF.36.5770), with two bacterial MscS channels from *E. coli* (WP_000389819) and *H. pilory* (WP_000343449.1). The position of the transmembrane domains TM1, TM2 and TM3 are underlined, the position of the putative gate residues are indicated by red arrows, conserved residues forming the hinges of TM3 are indicated with asterisks. The full alignment can be found in supplemental Figure S1. **B**. Arrangements of transmembrane helices of EcMscS (PDB ID 2OAU) and in the proposed homology model of TcMscS. The position of the putative gate and secondary constriction residues for TcMscS (F64 and L97) are indicated. **C.** Full homology TcMscS model in the closed and open states. N-terminal domain 1 (red), TM 2 (gold) and TM 3 (green) are followed by the cytoplasmic cage (cyan) and the C-terminal bundle (blue). The gate residues are indicated in pink and yellow.

The labeling of TcMscS helices in Fig. 1 matches the homologous segments of prokaryotic MscS. Because the TM1 segment in TcMscS is too short to cross the membrane, it is modeled in the conformation attached to the inner surface of the membrane (N1). The beta domain (cyan) is the last segment that aligns with EcMscS. The complete TcMscS homology model, which includes the bundled C-terminal segment, is shown in the predicted closed and open conformations (Fig. 1C). In addition to the F64 constriction (main gate), the beta domain that follows TM3 is predicted to bear a second hydrophobic constriction (Fig. 1C top panel, pink residues). This hypothetical C-terminal (cytoplasmic) gate formed by L97 (Fig. 1C, bottom panel, yellow residues) is narrower than the cage constriction present in *E. coli* MscS. The full model also shows the predicted opening transition that widens both gates. The overall displacement and tilting of TM3 helices on opening transition is modeled to increase the main gate radius from 3.9 to 6.5 Å accompanied by an in-plane area expansion of the protein by approximately 5.0 nm^2^. The *de novo* modeled flexible linkers lead to the C-terminal amphipathic helical domains hypothesized to form a coiled-coil bundle (blue), similar to what was observed in the crystal structures of MscL [59], another type of bacterial MS channel. The linkers and bundle might serve as a pre-filter to exclude larger osmolytes from occluding the pore. Overall, the structural model prediction supports the formation of a hepatmeric channel with a higher degree of conservation in the pore forming region and some key differences in the C-terminal segment, which is unique to TcMscS.

### Electrophysiological characteristics of TcMscS

To evaluate the functional properties of TcMscS, electrophysiological recordings were performed in *E. coli* giant spheroplasts (MJF641 Δ7 strain) expressing the channel. Single-channel currents were measured at low protein expression, with 1-3 channels per patch. Fig. 2A shows typical single-channel traces recorded under 110 mmHg pressure steps at +40 and −40 mV pipette voltages. The current amplitude of 12 pA translates to 0.40±0.02 nS conductance (n=11). The I-V curve for TcMscS obtained under symmetric electrolyte conditions (200 mM KCl) is linear (Fig 2B, black symbols), in contrast to EcMscS which shows a visible rectification [4, 60]. After a perfusion of buffer containing 425 mM KCl in the bath (Fig 2B, red symbols), the I-V curve changes its slope and the x-intercept reproducibly shifts to −4.3 ±0.4mV (n=4). According to the Goldman equation, under a given ionic gradient the reversal potential of −4.3 mV corresponds to a P_Cl-_/P_K+_ of 1.6, signifying that the channel passes about 3 Cl^-^ ions per 2 K^+^ ions, i.e., it shows a slight anionic preference.

**Figure 2.**
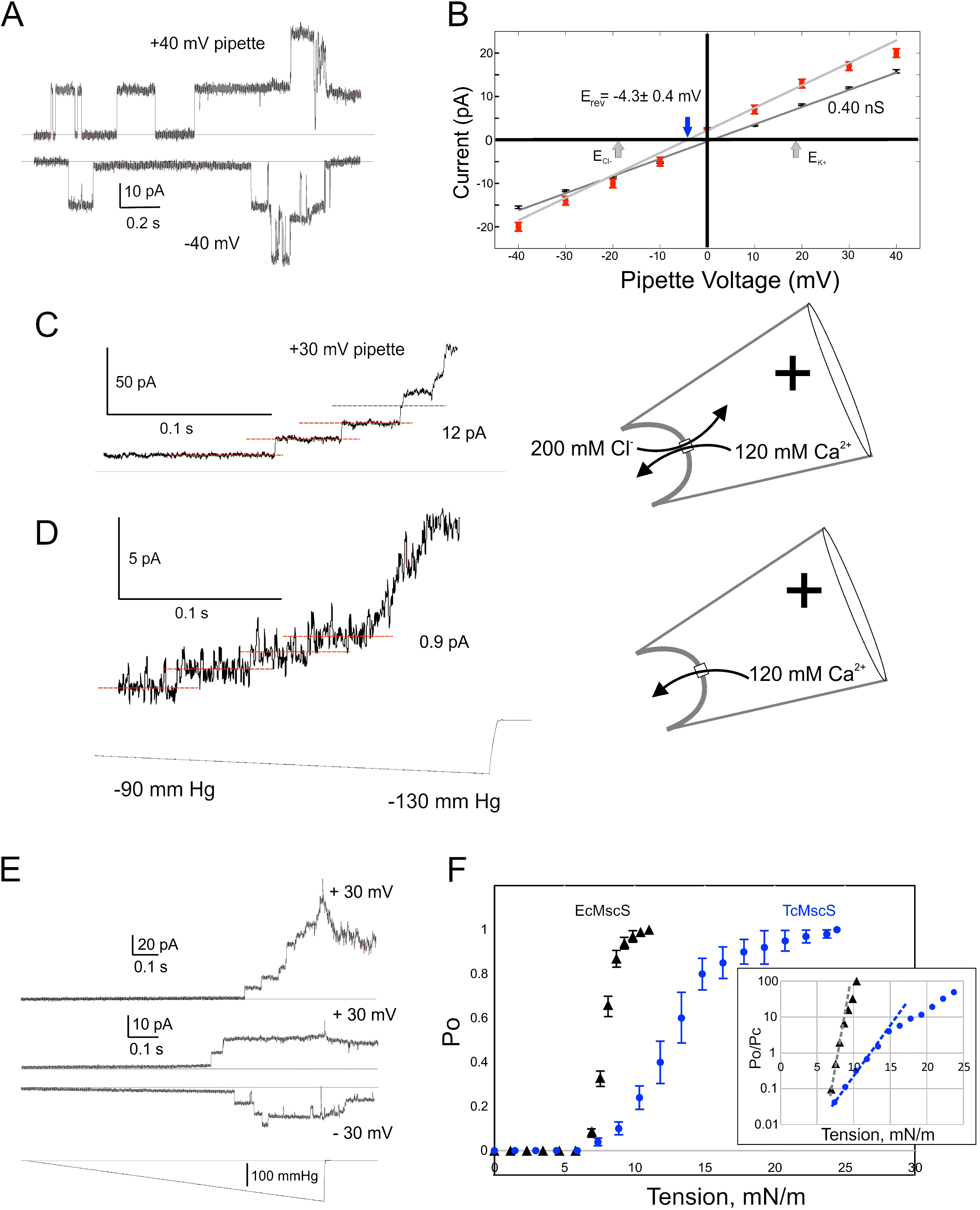
Electrophysiological properties of TcMscS. **A**. Single-channel currents at positive and negative pipette voltages **B.** Current-to-voltage relationships measured in symmetric (200 mM/200 mM KCl, black symbols, n=11) and asymmetric conditions (425mM/200 mM KCl, red symbols, n=4). The theoretical reversal potentials for Cl^-^ and K^+^ (grey arrows) and the experimental reversal potential measured for TcMscS (red arrow) are indicated. **C.** Unitary conductace of TcMscS recorded with 100 mM Ca gluconate, 20 mM CaCl_2_ in the pipette and 200 mM KCl in the bath at +30 mV in the pipette. **D.** The trace recorded from the same patch after perfusion of the bath with 100 mM Ca gluconate. In the latter configuration (shown on the right), Ca^2+^ is the dominant permeable ion. **E.** Examples of traces recorded with identical 1-s linear ramps on three separate patches bearing 11, 2 and 3 active channels at +30 mV or −30 mV pipette potential. **F.** Multiple ramp responses of TcMscS populations were normalized to the current at saturating pressure and converted to dose-response curves. The aggregated TcMscS curve (blue symbols, n=11) is compared with EcMscS dose-response curves (black symbols, n=54). The inset in panel **D** shows the same dose-response curves plotted as log (Po/Pc) and fitted with the Boltzmann equation. The shallower slope of the TcMscS dose-response curve indicates a smaller lateral expansion of the protein complex.

To explore whether the channel is permeable to Ca^2+^, we filled the pipette with 100 mM Ca gluconate (supplemented with 20 mM CaCl_2_ to ensure the silver chloride electrode works) and recorded unitary currents under asymmetric conditions with the standard 200 KCl buffer in the bath. Under positive pipette voltage (+30 mV), we recorded 12 ± 1 pA currents contributed by the inward flux of Ca^2+^ and outward flux of Cl^-^ (Fig. 2C). We then thoroughly perfused the bath solution with 100 mM Ca gluconate making Ca^2+^ the dominant permeable ion at this polarity. The amplitude of current dropped to 0.9 ±0.5 pA (n=3), translating into a 30 pS conductance (Fig. 2D). Our previous data [60] indicated that the unitary conductance of EcMscS scales linearly with the bulk conductivity of monovalent salt surrounding the channel. More recently, EcMscS was shown to be highly permeable to Ca^2+^ [7], its unitary conductance in 100 mM CaCl_2_ is about a half (0.5 nS) of the conductance in equimolar KCl (1.1 nS). In our experiment with TcMscS, the 400 pS unitary conductance in symmetric 200 mM KCl (24.1 mS/cm) drops to 30 pS in 120 mM Ca gluconate (4.64 mS/cm). The 5.2 fold drop in bulk conductivity resulted in a 13 fold decrease in unitary conductance, strongly suggesting that Ca^2+^ is still permeable through TcMscS, although at a lower level compared to monovalents. Interestingly, the TcMscS never opened at negative pipette potentials in the presence of high Ca^2+^.

Figure 2E shows two population currents recorded at + 30 or −30 mV in the pipette with identical ramp stimuli. The appearance of channels is essentially identical at two opposite voltages. The closing rate appears extremely slow as some channels remain open up to about 10 s after the stimulus ends.

To define the gating characteristics of TcMscS we analyzed the responses under pressure ramps and normalized the traces to saturating current to find the open probability and statistically assessed midpoint pressure required for activation (Fig. 2F). With EcMscS as a reference for known tension midpoint, we estimated the midpoint tension for TcMscS. EcMscS showed its half-activation pressure at 90 ± 3 mmHg (Fig. 2F, black symbols, n=54). TcMscS, had its pressure midpoint at 145 ± 20 mmHg (Fig. 2F, blue symbols n=11). Given that EcMscS activates in *E. coli* spheroplasts at 7.8 mN/m [61], we concluded that the midpoint of TcMscS activation is 12.6 mN/m. Although both channels start opening at approximately the same pressure (tension), the activation curve for TcMscS is shallower. Fitting the initial slopes of logarithmic Po/Pc plots (panel F, inset) using the two-state Boltzmann equation [62] gave estimates of the energies and in-plane area changes associated with gating. Gating of TcMscS was characterized by a 3 nm^2^ effective expansion of the protein, whereas the slope of EcMscS curve produced a 10.5 nm^2^ lateral expansion, consistent with models presented in Figure 1 and previous work by Anishkin et al [63].

These results unequivocally demonstrate that TcMscS is a mechanosensitive channel with functional characteristics comparable to the ones observed in bacterial MscS channels. It opens at sub-lytic tensions and, based on conductance, forms a ∼ 12 Å pore, which is essentially non-selective and capable of passing both anions and cations, including calcium, and possibly organic osmolytes such as betaine and other small amino acids.

### TcMscS expression and localization

To determine TcMscS localization, we generated polyclonal antibodies against the recombinant protein (*Supplementary materials and methods*). By immunofluorescence analysis (Fig. 3A) we found TcMscS mostly localized in the bladder of the contractile vacuole complex (CVC) of epimastigotes (Fig. 3A) and trypomastigotes (Fig. 3B). In epimastigotes, TcMscS colocalizes with VAMP7, another CVC protein [64, 65] (Fig. 4A), and partially colocalizes with actin (Fig. 4B). Interestingly, no colocalization with tubulin was observed (Fig. 4C). In trypomastigotes, TcMscS is found in the CVC bladder and is also distributed along the flagellum and the cell body (Fig. 3B), but shows no co-localization with the membrane marker SSP-1 (Fig. 4D). In intracellular amastigotes, TcMscS has a peripheral localization and no labeling was detected in the CVC (Fig. 3C). Colocalization with the amastigote membrane marker SSP-4 shows a clear overlap of the labeling (Fig.4E), indicating the translocation of TcMscS to the plasma membrane. The differential localization in the extracellular versus intracellular stages suggest that TcMscS could be playing different physiological roles in response to developmental and/or environmental cues. This hypothesis is reinforced by our data showing higher expression of TcMscS in epimastigotes compared with amastigotes and trypomastigotes, at the mRNA (Fig. 3D) and protein levels (Fig. 3E). Epimastigotes at the stationary phase of growth have a significantly lower amount of transcript compared with parasites in exponential growth (Fig. 3D), but no evident difference in the amount of protein was observed by Western blot analysis (data not shown).

**Figure 3.**
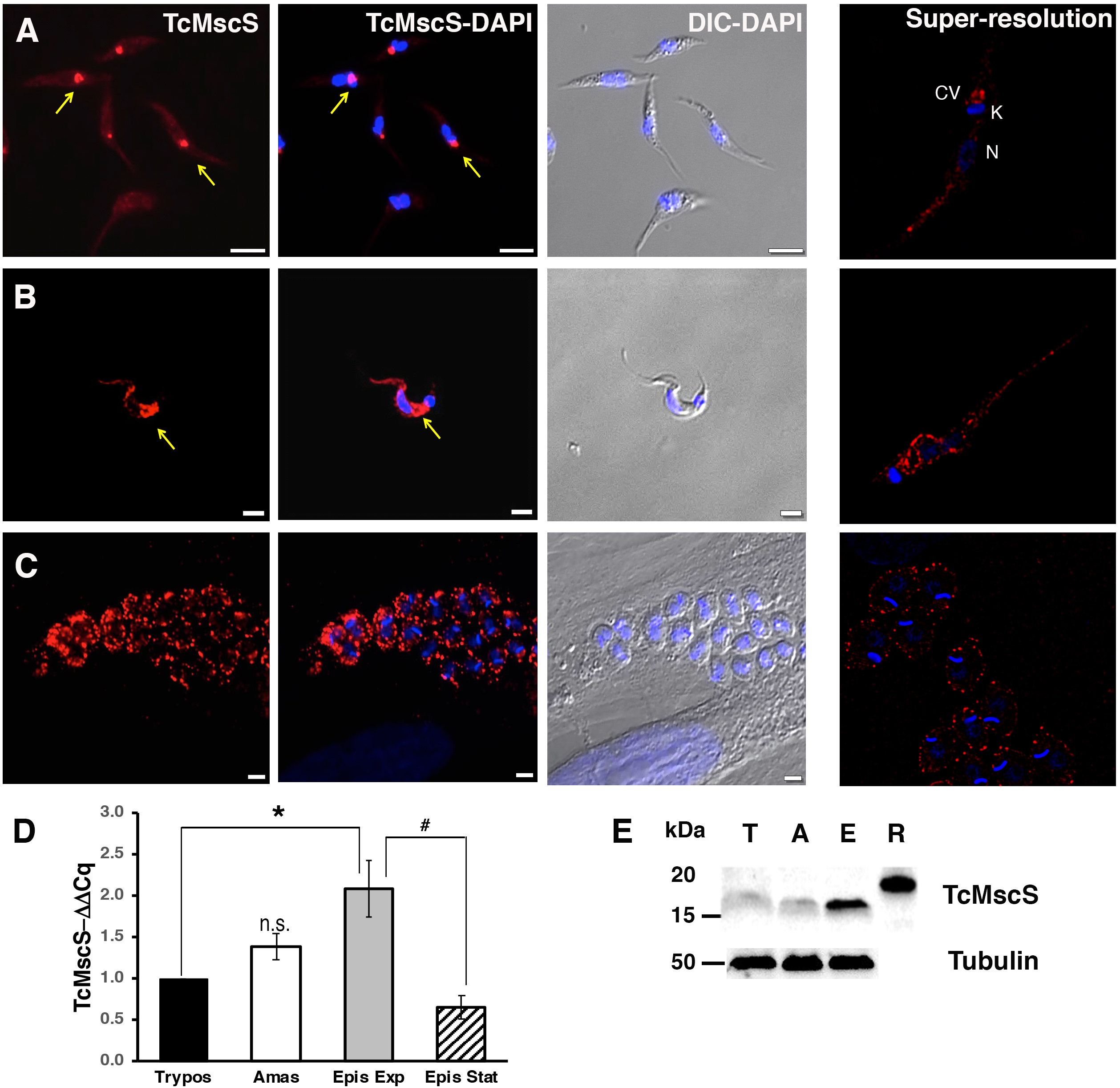
Localization and expression of TcMscS in *T. cruzi*. Immunolocalization of TcMscS (red) with specific polyclonal antibodies in epimastigotes **(A)**, trypomastigotes **(B)** and intracellular amastigotes **(C).** The right panels correspond to images of the same life stages obtained by super-resolution microscopy. Cellular structures are indicated as CV (contractile vacuole), K (kinetoplast) and N (nucleus). DNA was DAPI stained. Bar sizes: A = 5 µm; B and C = 5 µm. **D.** Expression of TcMscS in different life stages of the parasite, quantified by RT-qPCR. Trypomastigotes (Tryp) and amastigotes (Amast) were obtained from HEK-293 cells. Epimastigotes were analyzed at 4 days (Epis Exp) and 10 days of growth (Epis Stat). The values are indicated as ΔΔCq respect to the expression in trypomastigotes and normalized against GAPDH as housekeeping gene. The values are Mean ± SEM of 3 independent experiments in triplicate (*\* p = 0.006, # p = 0.009*). **E.** Western blot analysis of TcMscS in trypomastigotes (T), amastigotes (A) and epimastigotes (E). Purified recombinant protein (R) was used as positive control. The whole cell lysates were probed with polyclonal antibodies against TcMscS and monoclonal anti-tubulin was used as loading control.

**Figure 4.**
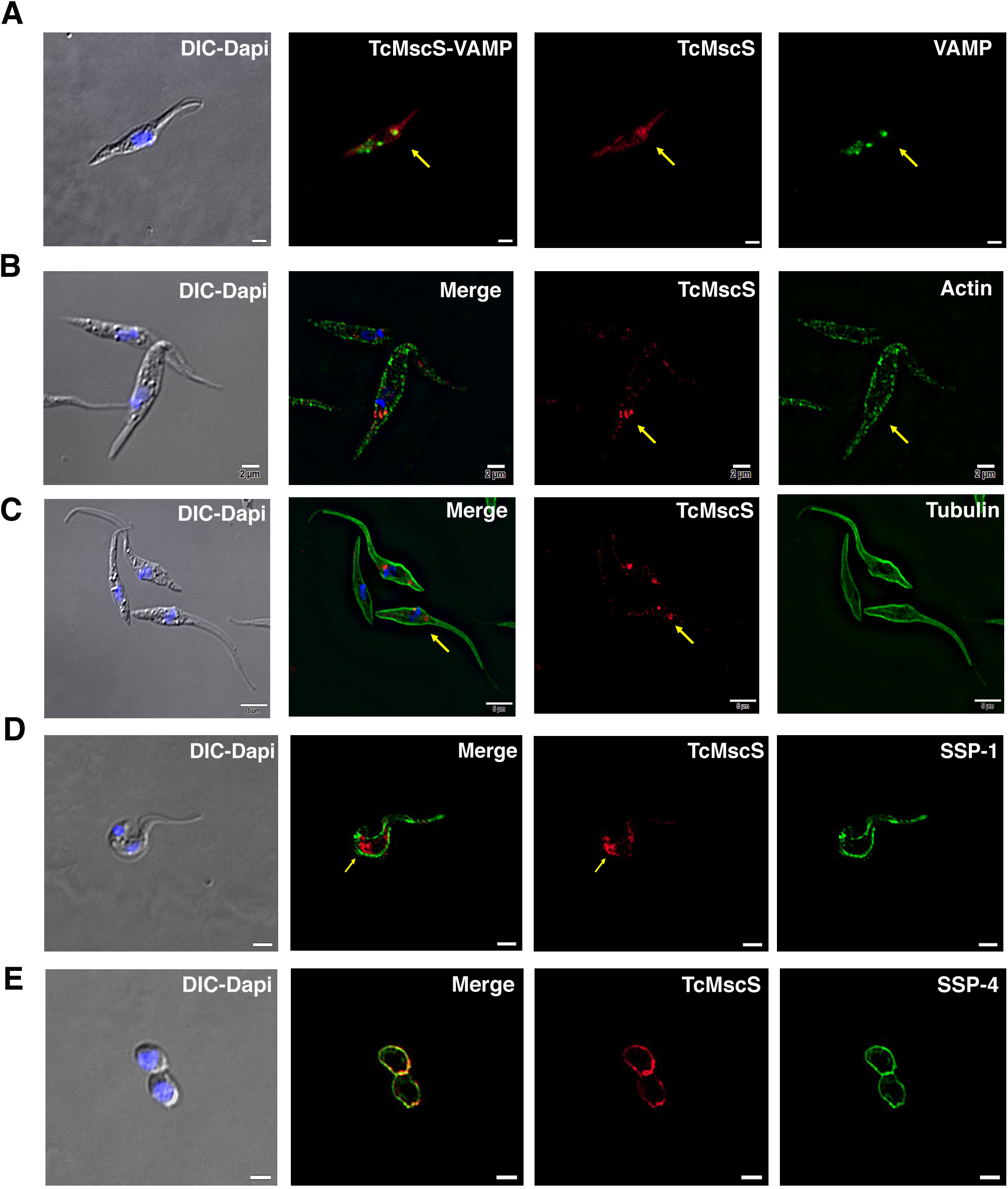
TcMscS colocalization with organelle markers. **A.** Immunofluorescence analysis of TcMscS (red) localization in epimastigotes overexpressing TcVAMP-GFP (green). **B.** Labeling of TcMscS (red) in WT Y strain epimastigotes partially colocalizes with anti-actin antibodies (green). **C.** Localization of TcMscS (red) and tubulin (green) in Y strain epimastigotes. No significant colocalization was observed. **D.** Immunolocalization of TcMscS (red) and membrane marker SSP-1 (green) in Y strain tissue-derived trypomastigotes. **E.** Co-localization of TcMscS (red) and membrane marker SSP-4 (green) in extracellular amastigotes obtained by host cell lysis. Nuclei and kinetoplasts were DAPI stained. Arrows indicate the position of the contractile vacuole. Bar size: 5 µm.

### TcMscS gene targeting by CRISPR-Cas9

To establish the physiological role of TcMscS in the parasites, we targeted the gene using the CRISPR-Cas9 system with (Fig. 5) or without DNA donor for gene replacement (Fig. S2). When we transfected the parasites with the Cas9/pTREX-n vector containing sgRNA2 (Table S1) targeting *TcMscS* without DNA donor, we observed a single nucleotide deletion (*del*177G), resulting in a frame shift and a premature stop codon (Fig 5A, top panel). These results were confirmed by genomic DNA sequencing in multiple independent samples. The truncated protein at amino acid 79 is not expected to be functional (Fig 5A, bottom panel) since the C-terminal end is absent. In these parasites, the transcript for TcMscS was still detected, although at significantly lower level compared with control parasites only expressing Cas9 and a scrambled sgRNA (Fig. 5C). Based on the level of mRNA and the detection of residual protein by Western-blot analysis (Fig. S3, A and B) it is possible that we are only targeting one of the alleles of the gene and we considered this strain a knockdown (TcMscS-KD). Importantly, the growth of TcMscS-KD epimastigotes is significantly reduced (Fig. S2 B, blue line) when compared to WT (Fig. S2 B, black line) and Cas9-scr controls (Fig. S2 B, red line). The parasites show abnormal morphology, with rounded bodies and formation of rosettes (Fig. S2 D, bottom panel).

**Figure 5.**
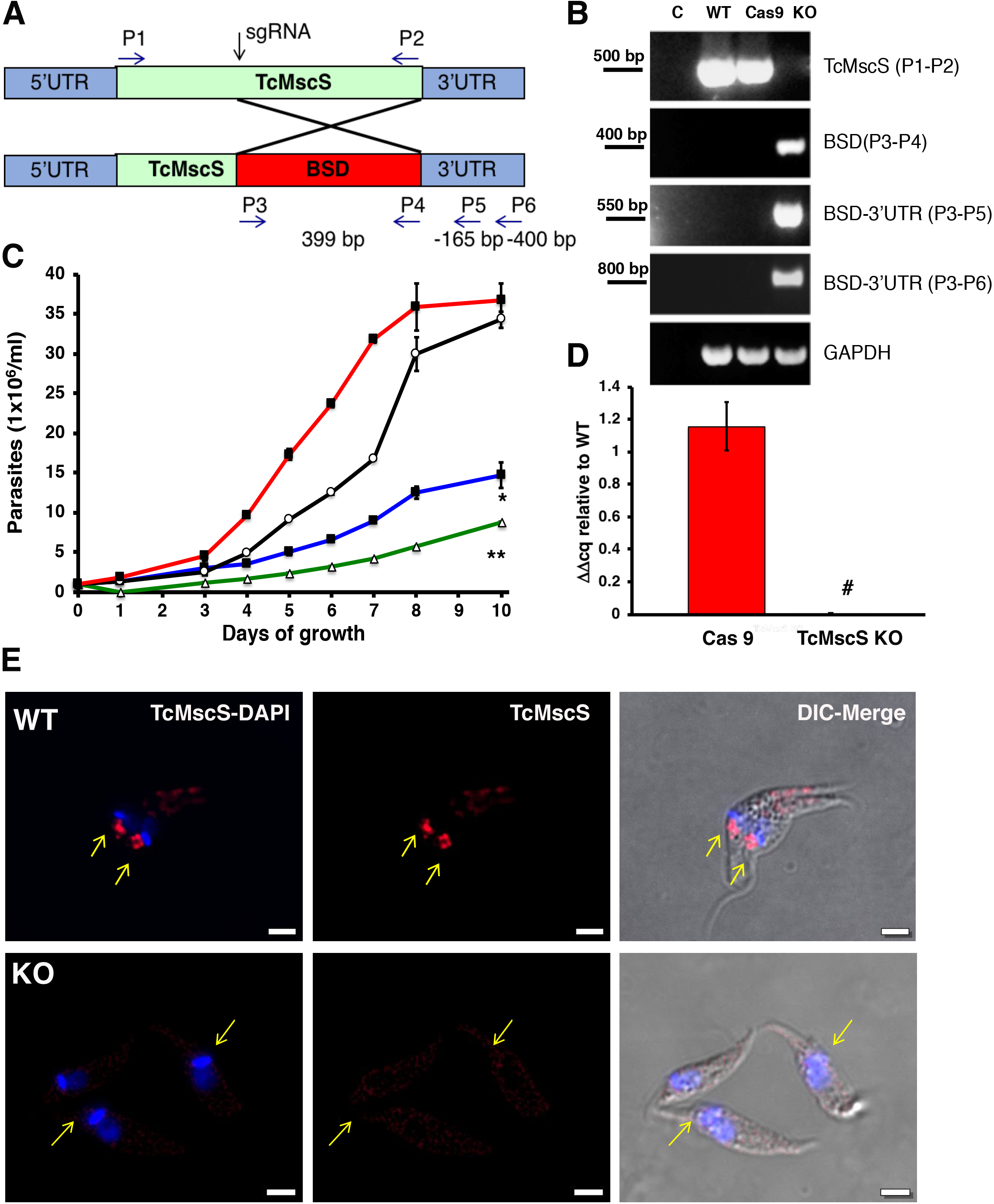
Phenotypic characterization of TcMscS KO. **A.** Schematic representation of *TcMcS* gene targeting by CRISPR/Cas9. A double strand DNA cut was induced at position +185 and repaired by homologous recombination with construct containing the *Bsd gene*. The primers used for screening are listed in Table S1. **B.** gDNA screening of TcMscS KO epimastigotes by PCR. Negative controls (C) without DNA are in lane 1 for all reactions. The ORF for *TcMscS* was amplified in WT and Cas9 controls but absent in TcMscS KO parasites. The correct insertion of the blasticidin cassette (BSD) was verified by amplification with primers annealing in ORF and the 3’UTR. GAPDH was used as a housekeeping gene for loading control. **C.** Growth curve of epimastigotes WT (black line), Cas9 (red line), TcMscS KD (blue line) and TcMscS KO (green line). All the experiments were started at 1×10^6^/ ml cells and counted daily. The values are Mean ± SEM of 3 independent experiments in triplicate (*, ** p<0.01). **D.** RT-qPCR quantifying the expression of TcMscS in epimastigotes control (Cas9) vs TcMscS-KO. The values are indicated as ΔΔCq respect to the expression in WT epimastigotes and normalized against GAPDH as housekeeping gene. All the samples were collected from parasites at 4 days of growth. The values are Mean ± SEM of 3 independent experiments in triplicate (*# p = 0.0001*). **E.** Immunofluorescence analysis of TcMscS (red) in epimastigotes Y strain (WT), TcMscS-KO. Nuclei and kinetoplasts were DAPI stained. The position of the contractile vacuole is indicated with yellow arrows. Bar size= 2 µm.

To completely abolish the expression of TcMscS, we co-transfected the parasites with sgRNA2-Cas9/pTREX-n vector plus a donor DNA cassette for homologous-directed repair (Fig. 5A). Genomic DNA analysis by PCR shows the absence of the *TcMscS* gene and the correct insertion of the blasticidin resistance cassette (Fig. 5B). No transcript was detected in epimastigotes (Fig. 5D) or trypomastigotes (Fig. S3, C). These results were confirmed by Western blot analysis (Fig. S3, A) and immunofluorescence (Fig. 5E), demonstrating the successful knockout of *TcMscS* (TcMscS-KO).

As expected, the growth rate of TcMscS-KO epimastigotes was significantly reduced (Fig. 5C, green line) and was lower than in TcMscS-KD parasites (Fig. 5C, blue line). The decrease in growth is accompanied by a marked reduction in motility (Suppl. Video 1 and 2), showing that the ablation of the channel has a negative effect on parasite fitness. To verify the specificity of the phenotype, we complemented the KO strains with the overexpression vector pTREX carrying a copy of TcMscS C-terminally tagged with myc. We successfully expressed the tagged construct (Fig. S4 A, C1 and C2), but as expected based on our previous data, the overexpression has a detrimental effect on the cells. This is in agreement with previous reports of porin and porin-like overexpression toxicity [66, 67]. The cell growth was not recovered (Fig. S4 B) and the epimastigotes showed abnormal morphology (Fig. S4 C). We then complemented the TcMscS-KO strain with a copy of the *T. brucei* ortholog TbMscS (Fig. S4 A and B, C3) and we were able to partially revert the cell growth phenotype, indicating some function conservation of the channel in other trypanosomatids.

### TcMscS regulates parasites osmotic stress responses

To evaluate the role of TcMscS in cell volume regulation we exposed the parasites to osmotic changes and followed the variations in absorbance over time as an indicator of variations in cell volume. Under hypoosmotic conditions of 115 mOsm/L (Fig. 6A) TcMscS-KD and KO epimastigotes experienced a maximum change in volume of about 38% compared with 23-25% in the controls (Table S2, Peak and Fig. 6B). When we analyzed the rate of recovery between 200-400 s, no significant difference was observed in the slope values, except for TcMscS-KO that had a slightly higher value. Nevertheless, KD and KO parasites do not regain their normal volume after 600 s, showing a remaining volume change of about 20% (Table S2, Final volume, Fig. 6C).

**Figure 6.**
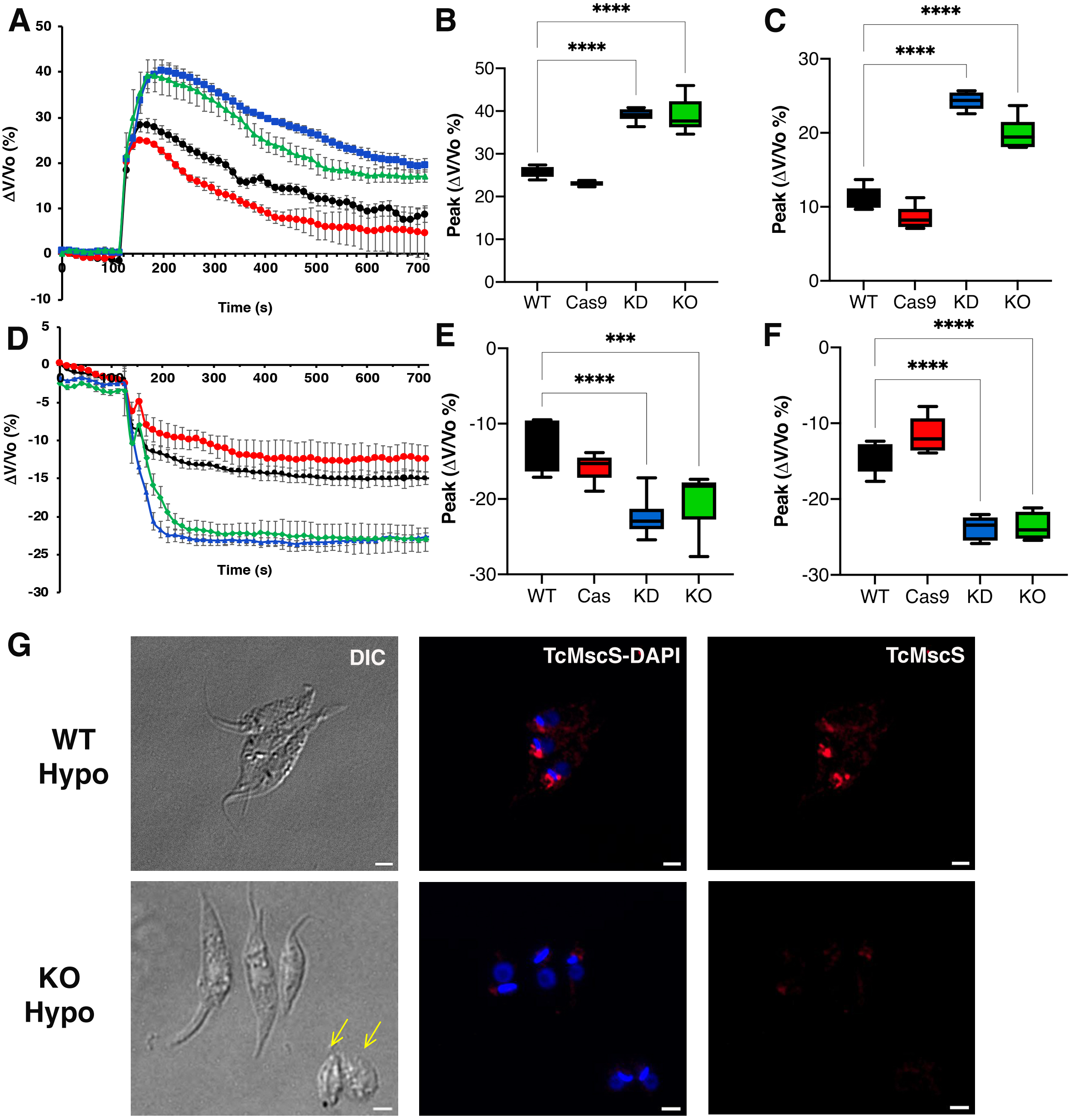
Osmotic stress responses in TcMscS mutants. **A.** Regulatory volume decreases in epimastigotes. Cells suspended in isosmotic buffer recorded for 120 sec and diluted with to a final osmolarity of 115 mOsm/L under constant ionic conditions. Relative changes in cell volume were monitored by determining absorbance at 550 nm over time, in WT (black), Cas9 (red), TcMscS-KD (blue) and TcMscS-KO (green). The absorbance values were normalized by the initial volume under isosmotic conditions and expressed as percentage of volume change. **B.** Analysis of the maximum volume change under hypoosmotic conditions. The peak area was calculated between 150-250 sec for all the experiments. **C.** Final volume recovery calculated between 600-700 sec. **D.** Volume decrease in epimastigotes after hyperosmotic shock. After 3 min under isosmotic conditions, cells were placed under hyperosmotic stress at 650 mOsm/L. Volume changes were monitored and calculated as explained in **A.** Peak analysis **(E)** and final volumes were calculated at the same times that in **B** and **C.** The values for A-F are mean ± SEM of 6 independent experiments in triplicate (p< 0.001). Statistical analysis details are listed in Table S2 and S3. **G.** Representative immunofluorescence images of WT and TcMscS-KO epimastigotes at 2 min post hypo-osmotic stress. TcMscS (red) was detected with specific polyclonal antibodies against the channel. Nuclei and kinetoplasts were DAPI stained. Yellow arrows indicate the presence of cells with abnormal morphology, observed in the TcMscS-KO strain. Bar size= 2 µm.

These results indicate that TcMscS, similar to other MscS channels, activates in response to increased membrane tension which results in redistribution of osmolytes and thus prevents excessive cell swelling. However, TcMscS-KD and KO parasites under hypoosmotic stress do not lose control of their volume completely, suggesting that TcMscS must act in conjunction with other mechanisms to orchestrate effective regulatory volume decrease responses.

Upon hyperosmotic stress of 650 mOsm/L (Fig. 6D), a similar phenotype was observed, with TcMscS-KD and TcMscS-KO parasite reducing their volumes by about 20%, compared with 12-15% in the controls (Table S3, Peak; Fig. 6E). As previously described [68], epimastigotes under hyperosmotic conditions do not regain normal volume, and parasites lacking TcMscS remained significantly more shrunken compared to controls (Fig. 6F), showing that TcMscS is required for regaining normal cell volume following hyperosmotic stress. We evaluated whether TcMscS localization changes upon osmotic stress and observed no differences in parasites under iso-, hypo- or hyperosmotic conditions at different time points (Fig. S5). It should be noted that TcMscS-KD and KO epimastigotes under isosmotic conditions show a cell volume about 36-38% higher than normal when compared with WT and Cas9 controls (Fig. S6). This significant difference underscores the role of TcMscS in the regulation of cell volume under normotonic conditions.

### TcMscS-KO causes cytosolic calcium dysregulation

TcMscS localization in the CVC and our electrophysiology data showing the permeation of calcium through the channel lead us to evaluate the cytosolic calcium level in the KO parasites. WT epimastigotes loaded with Fura 2-AM showed a baseline cytosolic Ca^2+^ of 102 ± 4 nM (n=9) and a robust increase upon addition of 1.8 mM extracellular CaCl_2_ (Fig. 7 A and B, black line) reaching 208 ± 8 nM 100 s post-stimulation. In contrast, TcMscS-KO had a lower baseline Ca^2+^ of 81 ± 4 nM (n= 8) and a lower increase post-addition (Fig. 7 A and B, green line). The increase in cytosolic free Ca^2+^ can be caused by release of the ion from intracellular stores and/or by uptake from the extracellular media. To evaluate whether the impairment of the KO was due to a reduced uptake, we treated the cells with Bay K8644, a known activator of voltage gated calcium channels (VGCC). These channels have been postulated as the main permeation pathway for extracellular Ca^2+^ in *T. cruzi* [69] but other reports have indicated its presence in the CVC [64]. When we stimulate the cells in the presence of Bay K (Fig. 7 C and D), WT parasites show a dramatic increase in cytosolic Ca^2+^, 10 fold higher than in non-pretreated cells (Fig. 7 A and C, black lines). TcMscS-KO exhibited an increase in cytosolic Ca^2+^ compared with non-Bay K conditions (Fig. 7 A and C, green lines), but the response was significantly lower than in WT (Fig. 7 D post-Bay K). Collectively, these results indicate the role of TcMscS in regulating the cytosolic Ca^2+^ dynamic on the parasites, probably by controlling intracellular stores such as the CVC. The lack of adequate Ca^2+^ homeostasis can severely impact cell motility and infectivity.

**Figure 7.**
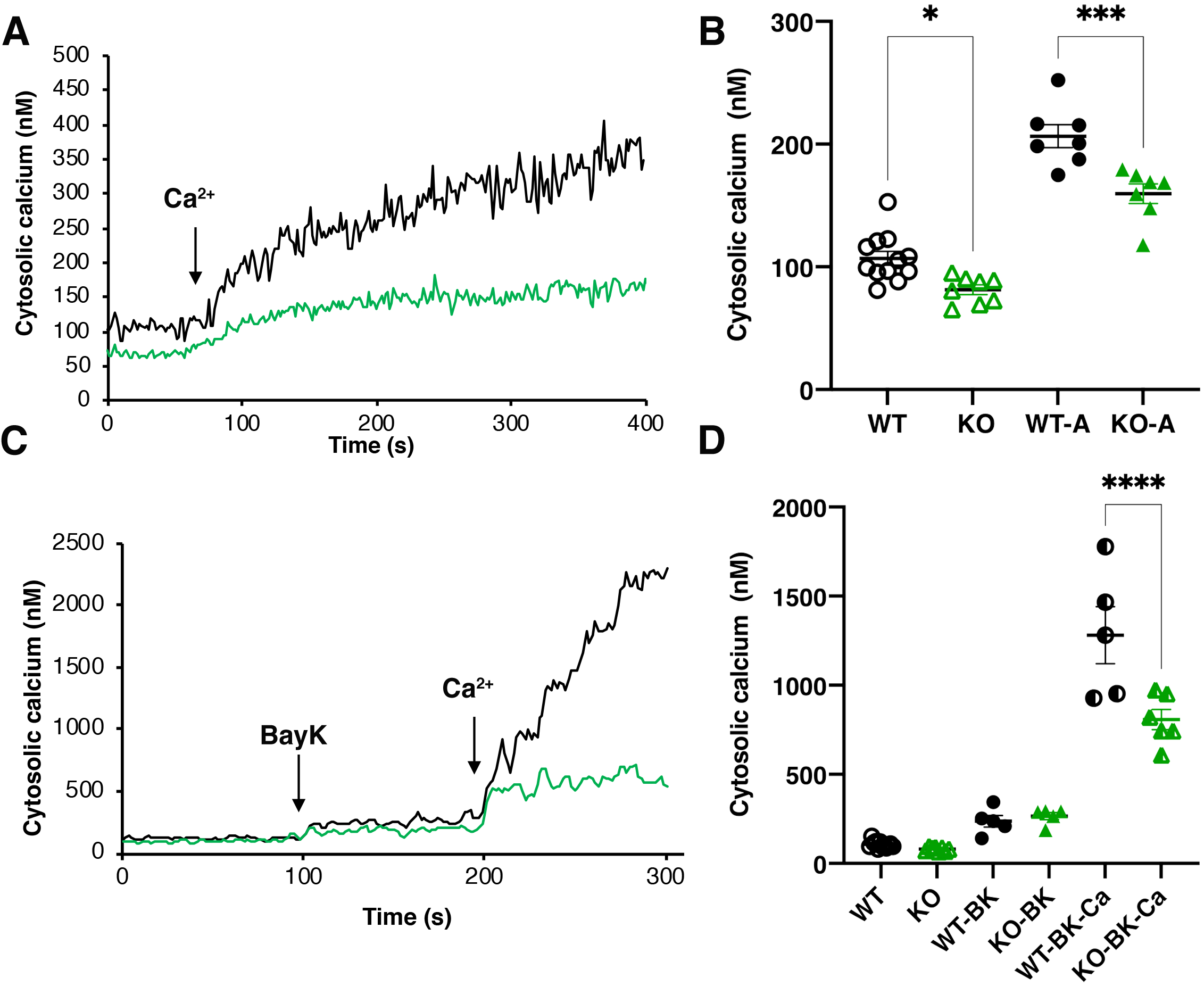
Intracellular calcium measurements. **A** and **C** are representative traces of intracellular calcium level in WT ( black line) or TcMscS-KO (green line) epimastigotes loaded with Fura 2-AM at baseline or after the addition of 1.8 mM CaCl_2_ and 10 μM of Bay K8644. **B** and **D** show the quantification of baseline Ca^2+^ concentration in the first 75 s and on the first 100 s after addition of the stimuli for WT (black) and TcMscS-KO (green) for multiple experiments. The values are Mean ± SE of n=7 independent experiments (*p<0.01, ***p<0.001, ****p< 0.0001).

### TcMscS-KO impairs parasite differentiation and infectivity

To elucidate the role of TcMscS in *T. cruzi’s* infectivity, we first evaluated the localization of the channel in metacyclic trypomastigotes (Fig. 8A). Similar to tissue-derived trypomastigotes, TcMscS is localized in the CVC bladder of the metacyclic forms, but also along the cell body, on the flagellar attachment zone region. The differentiation to metacyclic trypomastigotes *in vitro* is severely impaired in TcMscS-KO parasites, producing only 0.38% versus 6% obtained from WT epimastigotes after 72h of incubation in TAU media (Fig. 8B) [70]. We were able to infect mammalian cells with TcMscS-KO metacyclic trypomastigotes and obtained tissue culture derived trypomastigotes. The infective capacity of these parasites was not affected at early time points post-infection (Table S4) but did show a significant decrease in the production of intracellular amastigotes at 48 h post-infection (Fig. 8C and E; Table S4). These results confirm a defect in the capacity of the TcMscS-KO to replicate intracellularly. Quantification of extracellular trypomastigotes collected from the tissue culture supernatant shows a significant decrease in the number of TcMscS-KO parasites able to successfully differentiate into bloodstream trypomastigotes and egress from the cells (Fig. 8D). At 6 days post infection, TcMscS-KO produced a total of 5.2 x 10^6^ extracellular parasites, almost one order of magnitude lower than the amount produced by the Cas9-scr strain and WT parasites (Table S5).

**Figure 8.**
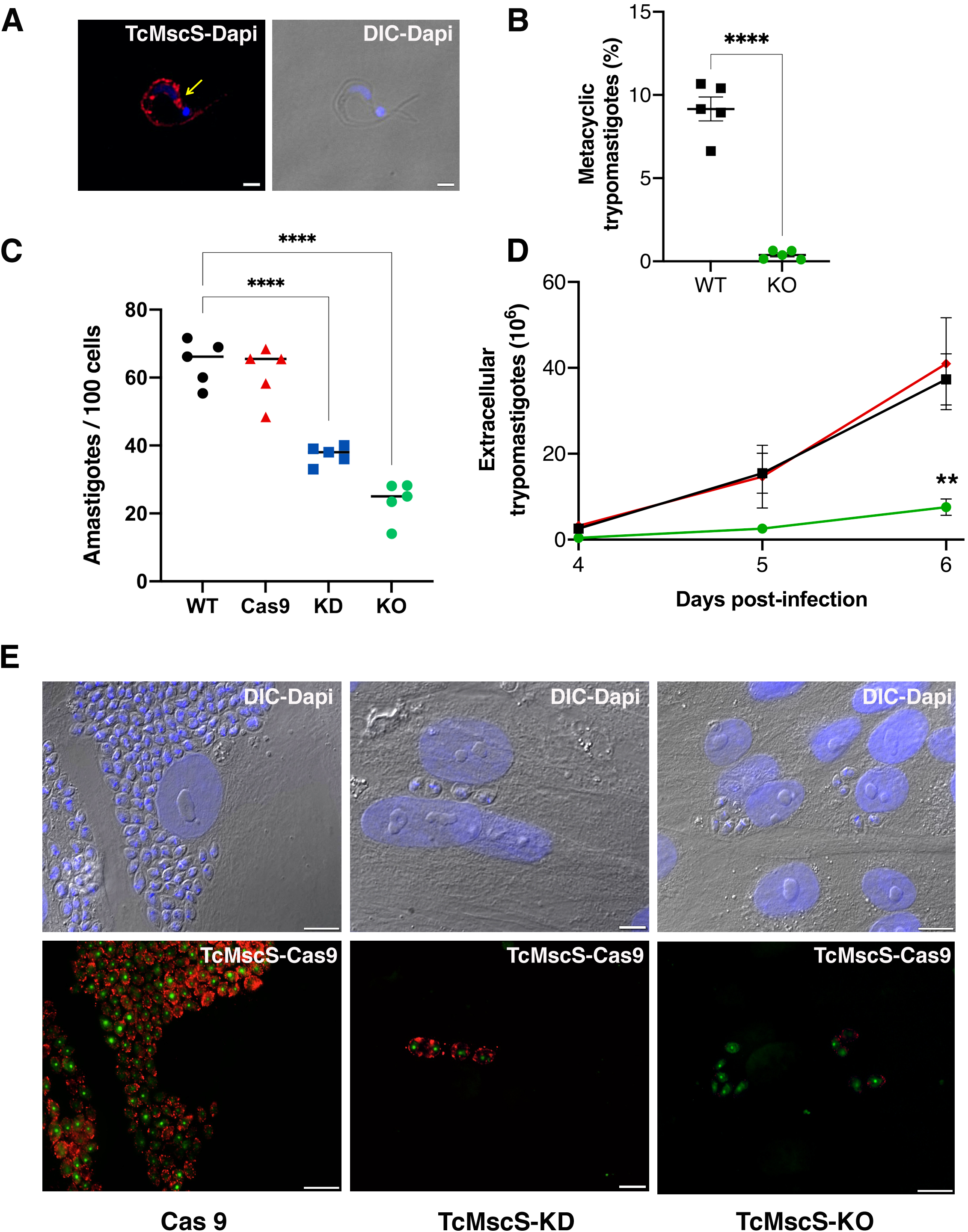
Infectivity defects in TcMscS mutants. **A.** Representative immunofluorescence images showing the localization of TcMscS in metacyclic trypomastigotes labeled with anti-TcMscS antibodies (red). The position of the contractile vacuole is indicated by yellow arrows. Nuclei and kinetoplasts were DAPI stained. Bar size= 2 µm. **B.** Percentage of metacyclic forms obtained *in vitro* from WT and TcMscS-KO parasites incubated for 72 h in TAU 3AAG media. Values are mean ± SEM of 5 independent experiments. **C.** Quantification of intracellular amastigotes at 48 h post infection. HEK 293 cells were infected at a MOI of 25:1 with trypomastigotes WT (black), Cas9 (red), TcMscS-KD (blue) or TcMscS-KO (green). TcMscS mutants show a significant decrease in intracellular amastigotes. Values are mean ± SEM of 5 independent experiments in triplicate. **D.** Quantification of extracellular trypomastigotes released from infected HEK293 cells after 4 days of infection. Cells were infected at a MOI of 25:1 with WT (black), Cas9 (red) or TcMscS-KO (green) trypomastigotes and extracellular parasites were collected daily for counting. Values are mean ± SEM of 4 independent experiments. **E.** Representative images of HEK293 cells infected with Cas9 (Cas9), TcMscS-KD or TcMscS-KO parasites. TcMscS was detected with specific antibodies (red), Cas9 was labeled with anti-GFP, nuclei and kinetoplasts were DAPI stained. Bar size= 10 µm. (**p<0.01, ****p< 0.0001).

In summary, the ablation of *TcMscS* results in loss of parasite fitness, impairing osmotic regulation, motility, calcium homeostasis, differentiation and cell infection capacity. The multiple phenotypic defects observed in TcMscS-KO parasites indicate that the physiological roles of this mechanosensitive channel have expanded beyond the primarily osmotic compensation reported for its bacterial ancestor EcMscS.

## Discussion

Propagation of *T. cruzi* involves alternating between mammalian and insect hosts; a life-history strategy that imposes significant mechanical demands on the parasite. This complex life cycle requires protective mechanisms and a strong differentiation potential that drives massive cell remodeling. The presence of TcMscS homologs in *Toxoplasma gondii*, *Plasmodium falciparum* and *Entamoeba histolytica* suggests a conserved function for these channels in other protozoan parasites [57]. Understanding their role as molecular transducers as well as likely effectors of mechanical cues throughout the parasites life cycle can provide new information about the role of physical and osmotic forces in the mechanisms regulating infectivity.

In this work we identify and present a functional characterization of the first MscS-like mechanosensitive channel in protozoan pathogens. TcMscS, a novel mechanosensitive channel found in *T. cruzi*, is required for growth, calcium homeostasis, differentiation and infectivity. The channel plays special roles in balancing osmotic and/or contractile forces in a stage- and location-specific manner. TcMscS belongs to the MscS superfamily and is structurally and functionally similar to the well characterized EcMscS [60]. MscS-like channels are a highly diverse group of proteins [71] found in Archaea [56], fungi [72–74], protozoans [75–77] and plants [51, 52, 78, 79], but not found in metazoans. The evolutionary origin of MscS-like channels can be traced to a common ancestor that was maintained across lineages as a protective mechanism against osmotic stress [80]. Beyond their primary role in volume regulation, these channels are also implicated in amino acid transport [81, 82] calcium homeostasis [73, 74, 83], and organelle biogenesis and maintenance [51-53, 78, 84, 85], highlighting the diversification of functions for this protein family over its evolutionary history [71, 86].

Expression in *E. coli* spheroplasts provided a clean experimental system to study activation by tension and single-channel properties of TcMscS. The linear non-rectifying conductance of the channel is approximately half of the conductance of EcMscS [60]. Similar to other MscS-like channels, TcMscS has a slight selectivity towards anions, but also shows measurable permability to calcium.

The size of the pore appears be sufficient for permeation of small organic osmolytes. In *T. cruzi*, regulatory volume decrease is driven primarily by efflux of amino acids and requires fluctuations in intracellular calcium [87]. Given the localization in the CVC and the non-selective nature of TcMscS, this channel could be a permeation pathway for amino acids and calcium required for cell volume regulation. An activation midpoint near 12 mN/m determined in bacterial spheroplasts suggests that the channel becomes fully active at tensions approaching the lytic limit, and thus it fulfills the role of a stress-releaving emergency valve, permitting the equilibration of osmolytes. TcMscS activities recorded in a heterologous expression system strongly suggest that the channel is gated directly by tension in the lipid bilayer without the need for cytoskeleton or other Tc-specific components. The ramp responses of TcMscS populations indicate a slow closing rate that allows the channel to remain open for a longer period, thus relieving the hypoosmotic stress more completely.

The channel has a novel 2TM topology (Fig. 1) and shows substantial homology with *E. coli* MscS only in the TM3 and beta domain region. Common to other tension-activated channels [88], TcMscS is predicted to have a short amphipathic N-terminal helix attached to the inner leaflet of the membrane. TcMscS has a completely unique C-terminal domain predicted to fulfill the function of pre-filter at the cytoplasmic entrance.

In most cell types, MscS channels are localized in the plasma membrane, but they are also found in internal organelles including the ER [73], contractile vacuole [50, 89–91] and plastids [53, 77]. In *T. cruzi*, TcMscS shows a differential localization in the extracellular versus intracellular stages. In epimastigotes, the channel has a defined intracellular localization in the bladder of the contractile vacuole and partially colocalizes with VAMP-7 and actin. There are no previous reports of actin associated to the bladder of the CVC, but we have shown the expression of myosin heavy chains V1 and V2 [64] suggesting the presence of a contractile machinery that might contribute to the periodic discharge of the organelle, similar to what has been described in *D. discoideum and Paramecium* [90, 92]. In tissue-derived bloodstream and *in vitro-*differentiated metacyclic trypomastigotes, TcMscS localization is expanded beyond the CVC to include the cell body and part of the flagellum. This suggests that the channel could be sensing changes in tension during flagellar beating.

The role of CVC as a key regulator of the cell volume in *T. cruzi* has been extensively characterized [47] and the presence of a mechanosensitive channel in CVC points to necessity of curbing excessive tensions to protect the organelle from osmotic lysis. Proteomic and functional analysis has shown the presence of numerous proteins involved in volume compensation [64], but no sensor of the osmotic state has been identified. We propose that TcMscS is activated upon excessive filling of the bladder, similar to what has been proposed in other protists [48, 92, 93]. If the normal exocytotic release of the bladder content fails to relieve pressure, the channel activation may release the content back to the cytoplasm to preserve the integrity of the CVC.

The peripheral localization of TcMscS in amastigotes deserves further consideration. Intracellular stages develop free in the cytosol under osmotically stable conditions, establishing persistent infection predominantly in muscle cells where contractility generates periodic stress [40]. It is known that *T. cruzi* infection causes extensive cytoskeletal remodelling and the amastigotes are not protected by a parasitophorous vacuole, thus, they can be mechanically perturbed [94–97]. At this stage, TcMscS main role could be detecting changes in tension and shear stress elicited by host cell contraction. Alternatively, these results can be interpreted considering a simple topological factor: the spherical amastigotes would be more sensitive to osmotic downshock, whereas elongated forms have room for swelling and are better protected by a stronger cytoskeleton, explaining the change of TcMscS localization in amastigotes. In addition, the channel may be facilitating cell volume adjustments in the course of active cell reshaping during differentiation to trypomastigotes, which are preparing to exit the cells.

TcMscS expression is developmentally regulated, with higher levels in epimastigotes at the exponential phase of growth. Changes in expression and localization of the channel could be responding to stage-specific cues, as well as external conditions, allowing the parasites to successfully transition through their developmental cycle. To test this hypothesis, we used CRISPR/Cas9 [98] to target TcMscS in the parasites and produced two strains with similar characteristics but different level of phenotypic defects. Neither the reduction (KD) or the elimination (KO) of the gene resulted in a lethal phenotype, but both cell lines showed a detectable increase in cell volume, a significant reduction in their extracellular and intracellular replication rates and impaired ability to produce metacyclic and bloodstream trypomastigotes, decreasing the infectivity of the parasites. The role of mechanical cues in differentiation has been demonstrated in several cell types including mesenchymal stem cells [11], osteoclasts [99] and neurons[14] where TRPV and PIEZO 1 mechanosensitive channels activity regulates developmental transitions [13, 100] and axonal growth [14]. Here, we provide the first evidence that mechanosensation mediated by TcMscS influences the developmental transition from non-infective to infective forms in *T. cruzi,* supporting the role of these channels in cell differentiation.

TcMscS-KO strains exhibited reduced motility and loss of infectivity. This is consistent with recent studies showing that motility is necessary for virulence in *T. brucei* [101]. In trypanosomatids, flagellar beating and motility are regulated by Ca^2+^ transients generated in a directional manner along the flagellum [102].The generation of Ca^2+^ waves linked to the activation of mechanosensitive channels has been demonstrated in *Chlamydomonas* [103], tubular kidney cells [104, 105], lung epithelial lining [106] and bone cells [11]. In *T. brucei*, a putative Ca^2+^ channel has been shown to be essential for flagellar attachment and cell viability [107] suggesting that Ca^2+^ influx is required for normal function. In *T. cruzi*, an ortholog of this Ca^2+^ channel has been identified in the CVC [64]. Our results show a dysregulation in the Ca^2+^ responses of TcMscS-KO, both at rest and after stimulation with calcium and Bay K. TcMscS seems to participate in the modulation of Ca^2+^ dynamics either directly, by regulating calcium release from intracellular stores like the CVC [48, 108] or indirectly by changing the ionic environment and membrane potential that control the activation of voltage-gated calcium channels. This association between the role of CVC in calcium regulation and TcMscS could explain the observed defects in motility and infectivity (Suppl. videos 1 and 2).

In yeast and plants MscS-like channels localized intracellularly regulate cytosolic Ca^2+^ oscillations required for organelle homeostasis and volume regulation [51, 73, 74, 77, 79, 84], supporting our hypothesis that TcMscS could be playing similar roles in *T. cruzi*.

As expected, the mutant parasites also showed a decreased ability to control cell volume both under stress and normal conditions, confirming the role of TcMscS in osmotic homeostasis. The compromised RVD observed in the knockout parasites suggest that other channels or transporters could be involved in the compensatory responses. This has been well documented in bacteria, where double knockout of MscS and MscL are required to elicit a defect in osmoregulation. In *T. cruzi*, bioinformatic analysis shows the presence of genes encoding for putative Piezo-like [57] and TRP-like channels[109]. Our RNAseq analysis of TcMscS-KO parasites do not show a significant change in the expression of those genes (data not shown). These results do not rule out the possibility of an increase in channel activity, independent of the mRNA expression level.

In summary, we have identified and functionally characterized the first mechanosensitive channel in trypanosomatids. TcMscS resembles bacterial MscS in its core, but lacks the first transmembrane helix and has a modified cytoplasmic domain. Beyond its canonical role in osmoregulation, the channel participates in calcium homeostasis and its ablation affects the capacity of the cells to proliferate, move, differentiate, and infect host cells. Our work opens new avenues to explore the connection between sensing of physical stimuli and the activation of signaling pathways implicated in differentiation and host-parasite interaction. It also identifies new effectors that bridge sensing of external cues and the factors that drive stage transitions of protozoans in a host-specific manner. Additionally, these types of channels are not present in mammalian cells, offering potential as novel and selective targets for drug development.

## Material and Methods

### Electrophysiological recordings of TcMscS

For electrophysiological studies in *E. coli* giant spheroplasts we used a N-terminally 6-His tagged version of *TcMscS* cloned into the expression plasmid pQE80 (Qiagen) with restriction sites BamHI and HindIII. TcMscS was expressed in the MJF641 (aka Δ7) strain devoid of seven endogenous MS channel genes [110]. The strain was a gift from Dr. Ian Booth (University of Aberdeen, UK). Competent MJF641 cells were transformed, selected overnight for transformants on plates with 100 g/ml ampicillin, and used immediately for spheroplast preparation. Spheroplast preparation was done following standard protocols [111]. Briefly, 3 ml of the colony-derived culture was transferred into 27 ml of LB containing 0.06 mg/ml cephalexin to block septation. After 1.5-2 hours of incubation, 1 mM IPTG was added to induce protein expression for 20 min. The filaments were transferred into a hypertonic buffer containing 1 M sucrose and subjected to digestion by lysozyme (0.2 mg/ml) in the presence of 5 mM EDTA resulting in spheres of 3-7 µm in diameter. The reaction was terminated by adding 20 mM Mg^2+^. Spheroplasts were separated from the rest of the reaction mixture by sedimentation through a one-step sucrose gradient.

### Recording conditions and protocols

Borosilicate glass (Drummond 2-000-100) pipets with tips 1-1.3 μm in diameter (bubble number 4.9-5.1) were used to form tight seals with the inner membrane. The tension-activated channel activities were recorded in excised inside-out patches. In most experiments the pipette and bath solution were symmetrical (200 mM KCI, 50 mM MgCI_2_, 5 mM CaCI_2_, 5 mM HEPES, pH 7.4). The bath solution was supplemented with 400 mM sucrose to osmotically balance the spheroplasts. An asymmetric configuration 425 mM /200 mM KCI (bath/pipette) was used to determine the channel’s anion/cation selectivity.

Calcium permeability was measured with a pipette solution containing 100 mM Ca gluconate, 20 mM CaCl2, 5 mM Hepes, pH 7.2 and bath solution containing only 100 mM Ca gluconate. The activities of TcMscS were recorded under a pressure ramp 0 to −200 mm Hg, at +30 mV or −30 mV voltage in the pipette. Traces were recorded using Clampex 10.3 software (MDS Analytical Technologies). Programmed mechanical stimuli were delivered using a modified high-speed pressure clamp apparatus (HSPC-1; ALA Scientific Instruments).

### Cell culture

Wild-type Y-strain (WT) epimastigotes were cultured in Liver Infusion Tryptose medium (LIT) as previously described [112]. Epimastigotes transfected with pTREX constructs including Cas9scr and TcMscS mutants (TcMscS-KD and TcMscS-KO) were cultured in LIT media supplemented with 10% FBS-HI and selection antibiotics (250 μg/mL of geneticin (G-418) and 15 μg/ml blasticidin) [98].

Tissue-derived bloodstream trypomastigotes and amastigotes were obtained as previously described [43]. Human foreskin fibroblast (HFF) and HEK-293 cells were cultured in High Glucose Dulbecco’s Modified Eagles Media (HG-DMEM) supplemented with 100 units/mL of penicillin, 100 μg/mL of streptomycin, 0.2% of amphotericin B, and 10% FBS-HI, and incubated at 37 °C with 5 % CO2.

### In Vitro Metacyclogenesis

Differentiation to metacyclic trypomastigote forms was induced under chemically defined conditions using triatomine artificial urine (TAU) medium as described [70]. Epimastigotes at 4 days of growth were collected by centrifugation at 1600 × g for 10 minutes, washed once in phosphate buffered saline solution (PBS) pH 7.4, resuspended in TAU media and incubated 2 hours at 28 °C. The supernatant was collected and resuspended in TAU with amino acids (TAU3AAG), incubated for up to 7 days at 28°C, collected by centrifugation resuspended in 5 mL of Dulbecco’s Modified Eagles Media (DMEM) supplemented with 20% fresh FBS to eliminate residual epimastigotes.

### In vitro infection assays

HEK-293 cells were plated onto coverlips in 12 well plates (1,000 cells/well) and incubated in supplemented HG-DMEM overnight at 37°C with 5% CO_2_. Infections were performed at a multiplicity of infection (MOI) of 25:1 with either WT, Cas9scr or TcMscS mutant trypomastigotes. After 6 hours, the cells were washed 3 times with Hank’s media and fresh DMEM was added. Coverslips were fixed in 4% paraformaldehyde-PBS at 6, 24 and 48 h, stained with DAPI (5 μg/ml) and mounted in Fluoromont media for quantification of intracellular parasites or processed for immunofluorescence. Immunofluorescence assays were performed to verify expression of Cas9-GFP and TcMscS in intracellular amastigote forms. Cells were incubated with anti-TcMscS (1:100) antibody and anti-GFP rabbit serum (Thermo Fisher Scientific, Inc.) (1:3,000) antibodies. Pictures were acquired in an Olympus IX83 microscope and processed with CellSens Dimension. All infection quantifications were done in 4 coverslips per experiment, in 3 or more independent experiments. At least 100 host cells were quantified per coverslip. The number of host cells vs. parasites was compared by a two-way ANOVA.

### Quantification of extracellular trypomastigotes

The amount of infective forms released from HEK-293 cells infected at a MOI of 25:1 was evaluated by collecting the supernatant of the cultures at days 4, 5 and 6 post-infection. To eliminate possible contamination with detached cells, the supernatant was first centrifuged at 300 x g for 5 minutes. The supernatant containing trypomastigotes was then centrifuged at 1,600 x g for 5 minutes and resuspended in 1 ml of DMEM. The cells were counted in a Z2 Beckman Coulter cell counter. All quantifications were done in triplicate for 3 independent experiments.

### TcMscS mutant generation by CRISPR/Cas9

Two strategies were developed to generate TcMscS-KD and TcMscS-KO mutants using the CRISPR/Cas9 one-vector system [98]. WT epimastigotes were transfected only with sgRNA/Cas9/pTREXn vector containing a sgRNA targeting *TcMscS*. Alternatively, the single vector containing TcMscS-sgRNA was co-transfected with a donor DNA to induce double strand break repair by homologous recombination, carrying a blasticidin resistance marker and 100 bp flanking regions of *TcMscS.* Three rotospacer adjacent motifs (PAM) sites within the *TcMscS* gene were identified and guide RNAs (sgRNA) were designed to specifically target the gene. To avoid off target effects, the protospacers were screened against the whole *T.* cruzi genome using ProtoMatch-V 1.0 kindly provided by Sebastian Lourido [113]. The sgRNAs targeting *TcMscS* were amplified with forward primers listed in Table S1 (P7, 8 and 9) and reverse P10 using plasmid pUC_sgRNA as template [98]. The amplicons were cloned into TOPO-Blunt vector and subcloned into sgRNA/Cas9/pTREXn vector. Successful cloning of TcMscS-sgRNAs was confirmed by sequencing. Two donor templates were constructed using the gene encoding blasticidin-S deaminase (Bsd) amplified from tdTomato/pTREX-b [98] and the ultramers indicated in Table S1 (P9-12). The first donor template contained 5’ and 3’-UTR regions of *TcMscS* flanking the *Bsd* cassette. The second donor template had the *Bsd gene* flanked by the last 100 bp of coding sequence of *TcMscS* and 100 bp of the 3’-UTR. Plasmid containing a scrambled sgRNA and Cas9 in a pTREXn vector were used as controls. To simplify the labeling the cell line expressing Cas9 and the scrambled sgRNA is labeled Cas9.

### Cell transfections

Y-strain epimastigotes were grown for 4 days under standard conditions, collected at 1600 × *g* for 10 minutes, washed once with BAG, and resuspended in 100 μL of Human T Cell Nucleofector Solution (Lonza). 5 × 10^7^ cells were mixed with 5 μg of each sgRNA plasmid with or without 5 μg of DNA donor and electroporated under the U-033 Amaxa Nucleofector protocol. Cells were allowed to recover overnight and appropriate antibiotics for selection (250 μg/mL G-418 and 15 μg/mL of blasticidin) were added.

### Genomic screening of transfectants

Clonal populations were obtained by serial dilutions, selected for 3 weeks and a total of 12 clones were screened for proper gene targeting or replacement. Total genomic DNA was extracted from WT, Cas9, TcMscS-KD and TcMscS-KO and analyzed by PCR to evaluate the presence of *TcMscS* (Table S1 P1-2) or the DNA donors (Table S1 P3-4 for blasticidin, P5-6 *TcMscS* downstream). GADPH amplification was used as a housekeeping gene for loading control (Table S1, P15-16). PCR results were confirmed by genomic DNA sequencing.

### Strain complementation

The complemented strains C1 and C2 were generated by introducing a copy of TcMscS where the PAM sequence downstream of the sgRNA was mutated by PCR with primers P17 and P18 (Table S1). The PAM-mutated ORF of *TcMscS* was amplified (primers P19-P20, Table S1) and subcloned into pTREX-p-myc with HindIII-XhoI restriction sites. The mutation was verified by sequencing and transfected into TcMscS-KO epimastigotes as described above.

Complemented strain C3 was obtained by overexpressing a copy of *T. brucei* homolog TbMscS. For that, Tb427.10.9030 sequence was amplified from genomic DNA extracted from WT procyclic forms 427 strain (purchased from BEI Resources) with primers P21-P22 (Table S1), and subcloned into pTREX-p-myc with restriction sites HindIII-XhoI. The complementation plasmid was transfected into TcMscS-KO epimastigotes and the parasites were selected with puromycin (2.5 μg/ml) for 5 weeks. The expression of the myc-tagged proteins was verified by western-blot with monoclonal anti-myc antibodies.

### TcMscS expression levels

TcMscS mRNA expression was assessed by RT-qPCR in the three life stages of the parasite as well as in WT, Cas9, TcMscS-KD and TcMscS-KO. Epimastigotes were collected during mid-log phase of growth, washed once with 1x PBS pH 7.4 and homogenized in TRI Reagent^®^. Total mRNA was extracted following the manufacturer protocol (Sigma-Aldrich, St. Louis, MO) followed by chloroform/ ethanol precipitation cDNA was obtained using SuperScript® III First-Strand Synthesis System (ThermoFisher Scientific, Inc. Waltham, MA) and oligo-dT_(20)_ primers. cDNA was analyzed by qPCR with Power SYBR Green PCR Master Mix (Applied Biosystems) and primers P1-P2 for *TcMscS.* Trypomastigotes were collected by centrifugation from supernatants of infected HEK-293 cells at 5 days post-infection. Amastigotes were collected from by mechanical lysis of infected HEK-293 cells at 3 days post-infections. RNA extraction and cDNA synthesis were performed as indicated for epimastigotes. All qPCR results were normalized against GAPDH as housekeeping gene (Table S1, P15-16) and indicated as ΔΔCq Mean ± SD of at least 3 independent experiments in triplicate.

### Growth curves

Growth of epimastigotes was measured in TcMscS-KD, TcMscS-KO, WT and Cas9 control parasites starting at a concentration of 1-5 × 10^6^ cells/mL in LIT media supplemented as previously described. The number of parasites was determined daily using a Z Series Coulter Counter (Beckman Coulter) The results are expressed as Mean ± SD of 3 independent experiments, in triplicate.

### Osmotic stress assays

Epimastigotes at log phase of growth were collected at 1,600 x *g* for 5 min, washed twice in PBS and resuspended in isosmotic buffer (64 mM NaCl, 4 mM KCl, 1.8 mM CaCl_2_, 0.53 mM MgCl_2_, 5.5 mM glucose, 150 mM D-mannitol, 5 mM Hepes-Na, pH 7.4, 282 mOsmol/L) at a cell density of 1 x 10^8^/ml. Aliquots of 100 μL of parasites were placed in a 96-well plate in isosmotic buffer and relative cell volume changes after osmotic stress were measured by light scattering method [114]. Briefly, pre-treatment readings of parasites in isosmotic buffer were taken for 3 minutes, followed by addition of 200 μL of hyposmotic buffer (64 mM NaCl, 4 mM KCl, 1.8 mM CaCl_2_, 0.53 mM MgCl_2_, 5.5 mM glucose, 5 mM HEPES-Na, pH 7.4) was added for a final osmolarity of 115 mOsmol/L. Alternatively, the cells were subjected to hyperosmotic stress with 200 μL of hyperosmotic buffer (64 mM NaCl, 4 mM KCl, 1.8 mM CaCl_2_, 0.53 mM MgCl_2_, 5.5 mM glucose, 500 mM D-mannitol, 5 mM HEPES-Na, pH 7.4). The absorbance at 550 nm was measured every 10 seconds for 12 minutes. Readings were normalized against the average of the three minutes of pre-treatment. Normalized absorbance readings were then converted into a percent volume change using the following equation: [(V_F_ - V_i_)/V_F_]× 100, where V_F_ is the absorbance value at the experimental time point and V_i_ is the initial absorbance value obtained at the same time point under isosmotic conditions.

The maximum volume change was calculated by peak function analysis between 150-250 sec. The final recovery volume was evaluated between 600-700 sec. Results are the Mean ± SE of 6 independent experiments in triplicate for all cell lines.

### Live video microscopy under osmotic stress

WT and TcMscS mutant epimastigotes were loosely adhered to poly-L-lysine treated glass bottom coverslips under isosmotic conditions. After adherence, the epimastigotes were subjected to hyposmotic stress under the same conditions described above. Live video microscopy was taken under DIC illumination for 600 seconds in an Olympus© IX83 inverted microscope system.

### Intracellular calcium measurements

Calcium measurements were done in WT (n=9) and TcMscS-KO (n=8) epimastigotes collected at four days of growth. Cells were pelleted at 1600g for 10 minutes at room temperature, washed three times with Buffer-A with Glucose (BAG: in mM NaCl 116, KCl 5.4, MgSO_4_ 0.8, glucose 5, HEPES 50 pH 7.3) and loaded with 5 µM Fura2-AM (Molecular Probes) in BAG for 30 minutes at 30℃, washed twice and resuspended in BAG at a concentration of 5×10^8^ cells/ml. Aliquots of 5×10^7^ cells were taken for each measurement with excitation at 340/380 nm and emission at 525 nm. Calibration was done by permeabilizing cells in BAG plus 1 mM EGTA and then adding increasing concentrations of CaCl_2_. The concentration of free calcium available was calculated using MaxChelator software (http://maxchelator.stanford.edu/) and the Kd was calculated according to the manufacturer protocol. Experimental recordings were allowed to stabilize at baseline before addition of 1.8 mM CaCl_2_ or 10 M of Bay K8644. Recordings were done on a Hitachi F7000 spectrofluorometer with excitation wavelength 340/380 nm and emission 510 nm.

### Statistical analysis

All the experiments were performed at least 3 times unless indicated otherwise, with intra-experiment replicates. The results are presented as Mean ± SD or SE as indicated in the figure legend. Statistical significance was evaluated by one way ANOVA with ad-hoc Bonferroni post-test or Student t-test as indicated in the figures.

## Acknowledgments

Dr. Roberto Docampo for his guidance and manuscript revisions, Stephen Vella and Dr. Myriam Hortua-Triana for assistance with super resolution imaging. Dr. Nikolas Nikolaidis kindly provided antibodies against actin.

This work was funded by the National Institute of Health NIAID grant R15AI122153 and American Heart Association grant 16GRNT30280014 to VJ, and scholarship from Cal State Fullerton MARC U*STAR Program grant 2T34GM008612-23 to JF.

## Notes

### Competing Interest Statement

The authors have declared no competing interest.

### Summary of Updates

Figures 1, 2, 5, 6 and 8 have been modified, Figure 7 contains new data, supplementary information has been reorganized and new methods incorporated.

