## Supplemental materials and figures for "A novel mechanosensitive channel controls osmoregulation, differentiation and infectivity in *Trypanosoma cruzi*"

**Table S1. Sequences of primers used for genetic constructs and PCR screening.** Bold letters indicate restriction sites. Highlighted sections are: in yellow protospacer sequences, in grey sections of ultramers that correspond to the flanking regions of the gene and in magenta 20 bp of the blasticidin resistance marker.

| Primer No. | Primer Name | Sequence |
|---|---|---|
| 1 | BamHI-TcMscS Forward | 5'-**GGATCC**ATGAAACGCTTTTTCAATCGCT-3' |
| 2 | HindIII-TcMscS Reverse | 5'- **AAGCTT**TCACTGCTTGGTTGCGTTGT-3' |
| 3 | Blasticidin Forward | 5'-ATGGCCAAGCCTTTGTCTCA-3' |
| 4 | Blasticidin Reverse | 5'-TTAGCCCTCCCACACATAAC-3' |
| 5 | -167 bp TcMscS downstream | 5'TGGCACGATCGGTGTCGATA-3' |
| 6 | -400bp TcMscS downstream | 5'**-** GTAATTTGTCCCTTCCTGGG-3' |
| 7 | sgRNA 1 Forward BamHI | 5'-GATC**GGATCC**GTAAGCGCTTTTCTGGTGCAGTTTTAGAGCTAGAAATAGC-3' |
| 8 | sgRNA 2 Forward BamHI | 5'-GATC**GGATCC**GGTGTAACAGGGGCCACAATGTTTTAGAGCTAGAAATAGC-3' |
| 9 | sgRNA 3 Forward BamHI | 5-GATC**GGATCC**GGTCAACAGTCAATTCGAACGTTTTAGAGCTAGAAATAGC-3' |
| 10 | sgRNA Reverse | 5'-CAGTGGATCCAAAAAAGCACCGACTCGGTG-3' |
| 11 | 100 bp 3'-end-Blasticidin Forward Ultramer | 5'AACATTTGTGATGAAACGCTTTTTCAATCGCTTCTATCTTGACACTGGCATTATTGCTGACCCCAGTCAACGTAGCCTCGCTAGTCGAGTAAGCGCTTTTATGGCCAAGCCTTTGTCTCA-3' |
| 12 | 100 bp of 3'-UTR-Blasticidin Reverse Ultramer | 5'TCACTGCTTGGTTGCGTTGTCTCCAGGTGTCTTGACAACGTCTTGTTTAGGGAGATCTGTTTCATGTGGGTTCTGCTTCTTTTCTTGTTCCCATGTCACTTTAGCCCTCCCACACATAAC-3' |
| 13 | 100 bp 5'-UTR-Blasticidin Forward Ultramer | 5'CGCTGCGCCTTTGCTTGTTGGCGTGTCGAACACTCTTTTTTTTTTTTTTTTTTTTGCTTTTTTTCTTTAACGCGTCGGTGAAGAGAGAGAAACATTTGTGATGGCCAAGCCTTTGTCTCA-3' |
| 14 | 3'-UTR-Blasticidin Reverse Ultramer | 5'CGGAGTTCAGAAAACCCAGCGAGACATGGTGAAACACAAGACGTTCTCAAACACATTCTTATTCCTCATTCCTTACTCCAGAAAAGTAAAGGTCACTCCCTTAGCCCTCCCACACATAAC-3' |
| 15 | GAPDH Forward | 5'- CAGAGCCTCAGTGTTGGTG-3' |
| 16 | GAPDH Reverse | 5'-TCAATACTCACTCTTGTTTGG-3' |
| 17 | PAM-MutationForw | 5'-CAGGGGCCACAATTGCATTTGCTTGCAAGGA-3' |

| | | |
|---|---|---|
| 18 | PAM-MutationRev | 5’-TCCTTGCAAGCAAATGCAATTGTGGCCCCTG-3’ |
| 19 | HindIII-TcMscS Forw | 5’-**AAGCTT**ATGAAACGCTTTTTCAATCGCT-3’ |
| 20 | Xho-TcMscSnonstopRev | 5’-CTTAGG**CTCGAG**CTGCTTGGTTGCGTTGTC-3’ |
| 21 | HindIII-TbMscS Forw | 5’-CTTAGG**AAGCTT**ATGAAGCGATTTTTTGACAAG-3’ |
| 22 | Xho-TbMscSnonstopRev | 5’-CTTAGG**CTTGAG**CCTGACTTTACTTCGTTC-3’ |

**Table S2: Changes in cell volume upon hypoosmotic stress**

| | WT | Cas9 | TcMscS-KD | TcMscS-KO |
|---|---|---|---|---|
| Peak (%) | 25.83±0.52 | 23.08±0.23 | 39.10±0.63* | 38.96±1.64* |
| p-value | | 0.575 | $2.43E^{-09}$ | $8.65E^{-05}$ |
| Recovery slope | -0.0360±0.0029 | -0.0327±0.0057 | -0.0471±0.0031 | -0.0685±0.002* |
| p-value | | 0.629 | 0.025 | $7.3E^{-06}$ |
| Final volume | 8.86±0.71 | 7.99±2.05 | 20.48±0.65* | 17.60±0.68* |
| p-value | | 0.7099 | $2.96E^{-07}$ | $4.95E^{-06}$ |

For all the conditions values are Mean±SE of n=5- p values were calculated based on one-way ANOVA analysis with Bonferroni post-test. Differences were considered significant when p<0.01(*).

**Table S3: Changes in cell volume upon hyperosmotic stress**

| | WT | Cas9 | TcMscS-KD | TcMscS-KO |
|---|---|---|---|---|
| Peak (%) | -12.41±1.36 | -15.78±0.74 | -22.43±1.12* | -20.10±1.59* |
| p-value | | 0.0055 | 0.0002 | 0.0043 |
| Final volume | -14.38±0.93 | -11.59±1.08 | -23.84±0.71* | -23.56±0.82* |
| p-value | | 0.087 | $4.27E^{-05}$ | $7.99E^{-05}$ |

For all the conditions values are Mean±SE of n=5. p values were calculated based on one-way ANOVA analysis with Bonferroni post-test. Differences were considered significant when p<0.01(*).

**Table S4: Quantification of intracellular amastigotes**

| Amas/100 cells | WT | Cas9 | TcMscS-KD | TcMscS-KO |
|---|---|---|---|---|
| Average 6 h-pi | 2.49±0.90 | 1.41±0.65 | 3.46±0.96 | 3.09±1.26 |
| p-value | | 0.36 | 0.48 | 0.71 |
| Average 48 h-pi | 56.43±5.73 | 54.96±5.20 | 37.12±1.17* | 25.64±1.06* |
| p-value | | 0.85 | 0.029 | 0.006 |

For all the conditions values are Mean±SE of n=5. p values were calculated based on one-way ANOVA analysis with Bonferroni post-test. Differences were considered significant when $p<0.05$(*).

**Table S5: Quantification of extracellular trypomastigotes**

| Trypomastigotes ($10^6$) | WT | Cas9 | TcMscS-KO |
|---|---|---|---|
| 4 dpi | 2.11±0.74 | 4.04±1.09 | 0.41±0.03 |
| 5 dpi | 12.6±4.34 | 14.2±5.19 | 1.83±0.48 |
| 6 dpi | 31.8±6.91 | 49.3±11.2 | 5.22±1.27* |
| p-value (6 dpi) | | 0.24 | 0.003 |

For all the conditions values are Mean±SE of n=4. p values were calculated based on one-way ANOVA analysis with Bonferroni post-test. Differences were considered significant when $p<0.05$(*).

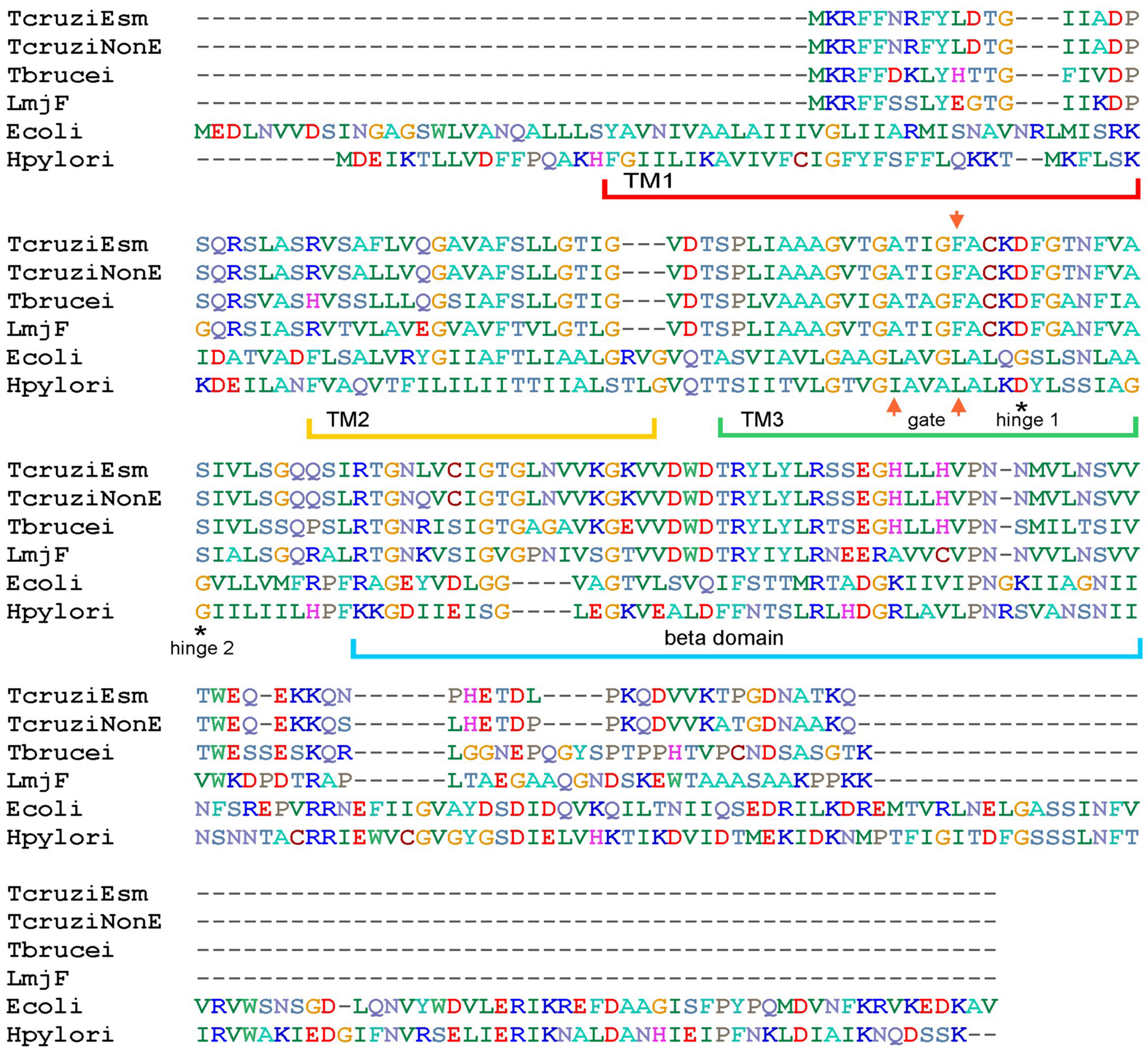
TcruziEsm ---------------------------------------MKRFFNRFYLDTG---IIADP
TcruziNonE ---------------------------------------MKRFFNRFYLDTG---IIADP
Tbrucei ---------------------------------------MKRFFDKLYHTTG---FIVDP
LmjF ---------------------------------------MKRFFSSLYEGTG---IIKDP
Ecoli MEDLNVVDSINGAGSWLVANQALLLSYAVNIVAALAIIIVGLIIARMISNAVNRLMISRK
Hpylori ---------MDEIKTLLVDFFPQAKHFGIILIKAVIVFCIGFYFSFFLQKKT--MKFLSK
TM1
TcruziEsm SQRSLASRVSAFLVQGAVAFSLLGTIG---VDTSPLIAAAGVTGATIGFACKDFGTNFVA
TcruziNonE SQRSLASRVSALLVQGAVAFSLLGTIG---VDTSPLIAAAGVTGATIGFACKDFGTNFVA
Tbrucei SQRSVASHVSSLLLQGSIAFSLLGTIG---VDTSPLVAAAGVIGATAGFACKDFGANFIA
LmjF GQRSIASRVTVLAVEGVAVFTVLGTLG---VDTSPLIAAAGVTGATIGFACKDFGANFVA
Ecoli IDATVADFLSALVRYGIIAFTLIAALGRVGVQTASVIAVLGAAGLAVGLALQGSLSNLAA
Hpylori KDEILANFVAQVTFILILIITTIIALSTLGVQTTSIITVLGTVGIAVALALKDYLSSIAG
TM2
TM3
gate
*
hinge 1
TcruziEsm SIVLSGQQSIRTGNLVCIGTGLNVVKGKVVDWDTRYLYLRSSEGHLLHVPN-NMVLNSVV
TcruziNonE SIVLSGQQSLRTGNQVCIGTGLNVVKGKVVDWDTRYLYLRSSEGHLLHVPN-NMVLNSVV
Tbrucei SIVLSSQPSLRTGNRISIGTGAGAVKGEVVDWDTRYLYLRTSEGHLLHVPN-SMILTSIV
LmjF SIALSGQRALRTGNKVSIGVGPNIVSGTVVDWDTRYIYLRNEERAVVCVPN-NVVLNSVV
Ecoli GVLLVMFRPFRAGEYVDLGG----VAGTVLSVQIFSTTMRTADGKIIVIPNGKIIAGNII
Hpylori GIILIILHPFKKGDIIEISG----LEGKVEALDFFNTSLRLHDGRLAVLPNRSVANSNII
*
hinge 2
beta domain
TcruziEsm TWEQ-EKKQN------PHETDL----PKQDVVKTPGDNATKQ------------------
TcruziNonE TWEQ-EKKQS------LHETDP----PKQDVVKATGDNAAKQ------------------
Tbrucei TWESSESKQR------LGGNEPQGYSPTPPHTVPCNDSASGTK-----------------
LmjF VWKDPDTRAP------LTAEGAAQGNDSKEWTAAASAAKPPKK-----------------
Ecoli NFSREPVRRNEFIIGVAYDSDIDQVKQILTNIIQSEDRILKDREMTVRLNELGASSINFV
Hpylori NSNNTACRRIEWVCGVGYGSDIELVHKTIKDVIDTMEKIDKNMPTFIGITDFGSSSLNFT
TcruziEsm ---------------------------------------------------
TcruziNonE ---------------------------------------------------
Tbrucei ---------------------------------------------------
LmjF ---------------------------------------------------
Ecoli VRVWSNSGD-LQNVYWDVLERIKREFDAAGISFPYPQMDVNFKRVKEDKAV
Hpylori IRVWAKIEDGIFNVRSELIERIKNALDANHIEIPFNKLDIAIKNQDSSK--

Fig. S1

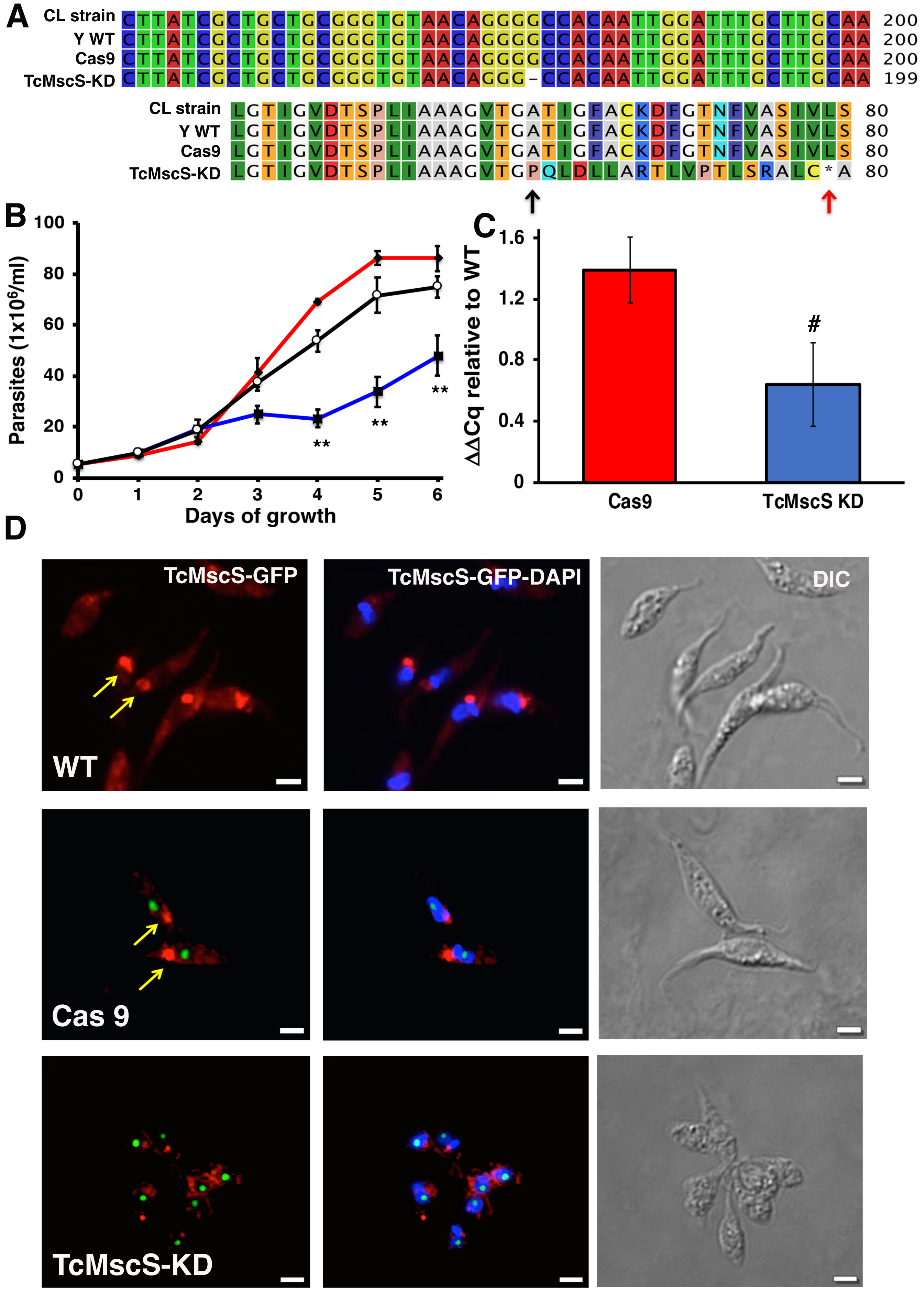
A
CL strain CTTATCGCTGCTGCGGGTGTAACAGGGGCCACAATTGGATTTGCTTGCAA 200
Y WT CTTATCGCTGCTGCGGGTGTAACAGGGGCCACAATTGGATTTGCTTGCAA 200
Cas9 CTTATCGCTGCTGCGGGTGTAACAGGGGCCACAATTGGATTTGCTTGCAA 200
TcMscS-KD CTTATCGCTGCTGCGGGTGTAACAGGG-CCACAATTGGATTTGCTTGCAA 199
CL strain LGTIGVDTSPLIAAAGVTGATIGFACKDFGTNFVASIVLS 80
Y WT LGTIGVDTSPLIAAAGVTGATIGFACKDFGTNFVASIVLS 80
Cas9 LGTIGVDTSPLIAAAGVTGATIGFACKDFGTNFVASIVLS 80
TcMscS-KD LGTIGVDTSPLIAAAGVTGPQLDLLARTLVPTLSRALC*A 80
B
Parasites (1x10⁶/ml)
100
80
60
40
20
0
**
**
**
0 1 2 3 4 5 6
Days of growth
C
ΔΔCq relative to WT
1.6
1.2
0.8
0.4
0
#
Cas9
TcMscS KD
D
TcMscS-GFP
TcMscS-GFP-DAPI
DIC
WT
Cas 9
TcMscS-KD

Fig. S2

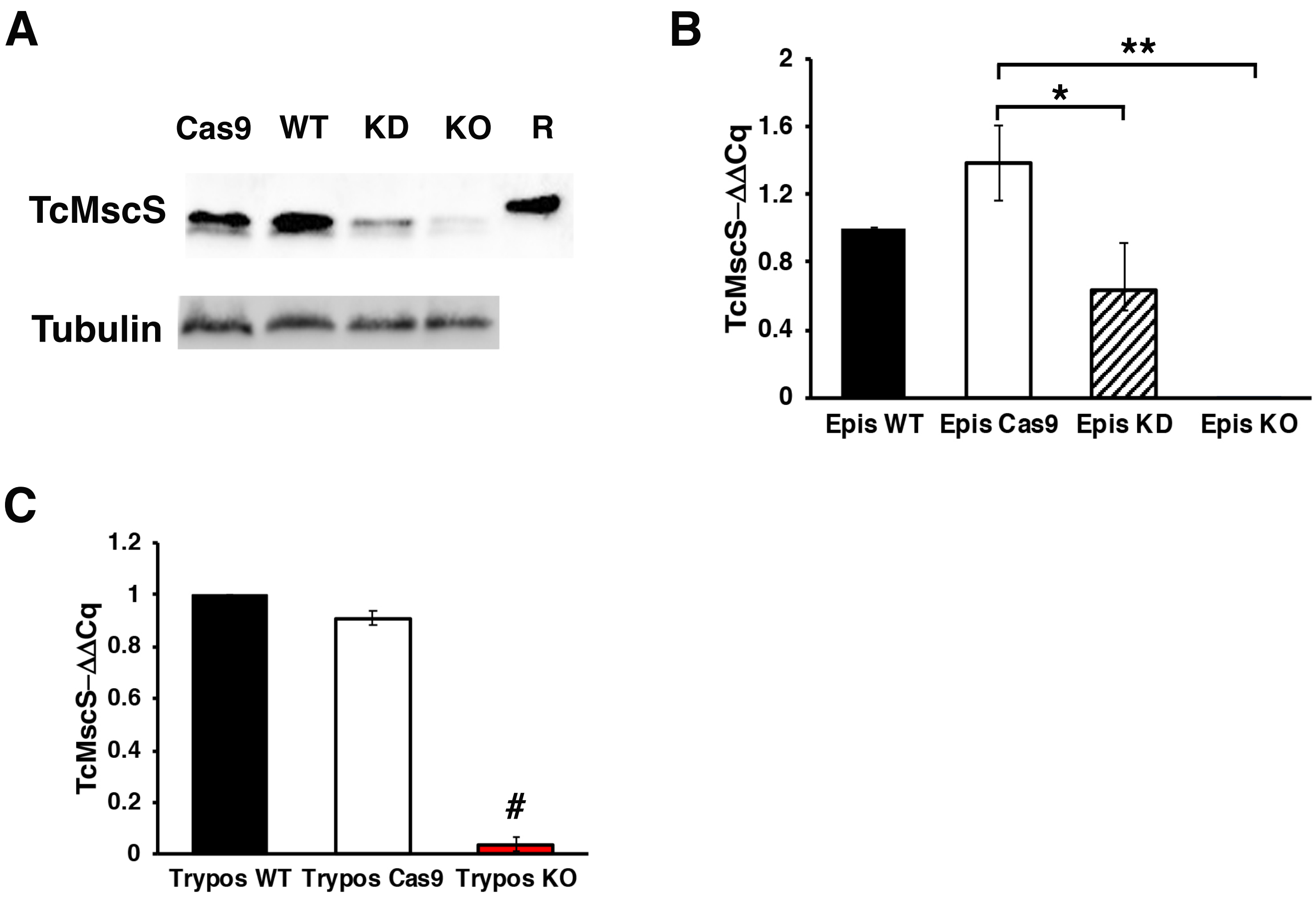
A
Cas9
WT
KD
KO
R
TcMscS
Tubulin
B
TcMscS−ΔΔCq
2
1.6
1.2
0.8
0.4
0
**
*
Epis WT
Epis Cas9
Epis KD
Epis KO
C
TcMscS−ΔΔCq
1.2
1
0.8
0.6
0.4
0.2
0
#
Trypos WT
Trypos Cas9
Trypos KO

Fig. S3

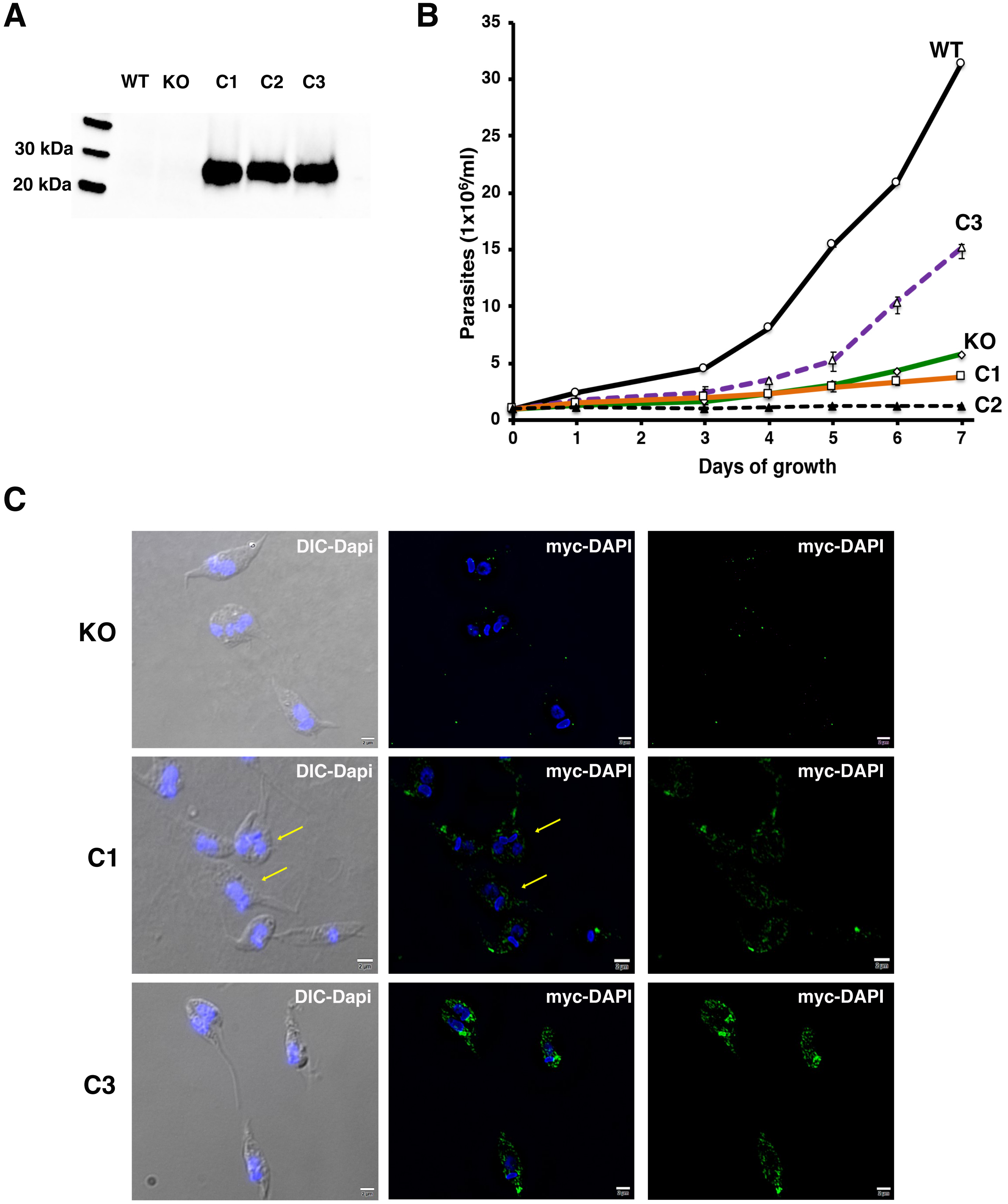
A
WT
KO
C1
C2
C3
30 kDa
20 kDa
B
Parasites (1x10⁶/ml)
Days of growth
0
5
10
15
20
25
30
35
0
1
2
3
4
5
6
7
WT
C3
KO
C1
C2
C
KO
C1
C3
DIC-Dapi
myc-DAPI
myc-DAPI

Fig.S4

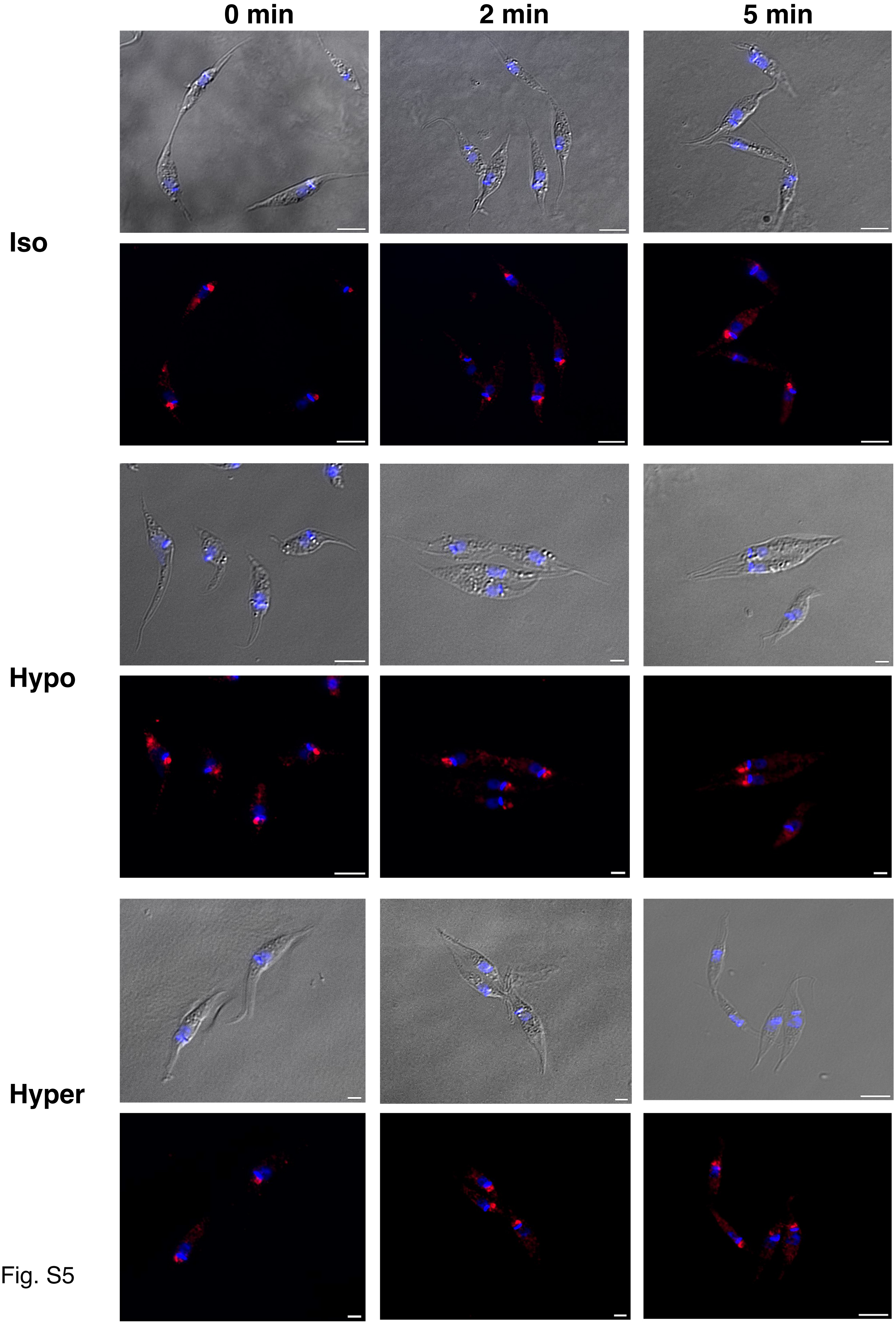


Fig. S5

**A**

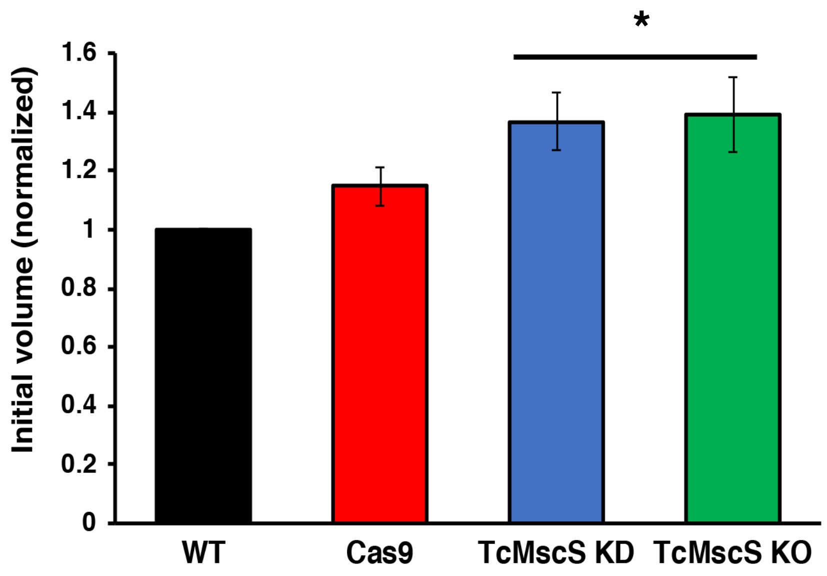
*
1.6
1.4
1.2
1
0.8
0.6
0.4
0.2
0
Initial volume (normalized)
WT
Cas9
TcMscS KD
TcMscS KO

Fig. S6

**Supplemental material - Dave et al.**

### 1) Supplemental materials and methods

*TcMscS sequence analysis*

Predicted protein sequences for TcMscS CL Brener Esmeraldo-like (TcCLB.504171.40) and Non-Esmeraldo-like haplotype (TcCLB.509795.40) were compared with the putative sequences for *Trypanosoma brucei* (Tb427.10.9030) and *Leishmania major* (LmjF.36.5770) MscS-like channels. Multiple alignments were compared with sequences for *E. coli* (WP_000389819) and *H. pilory* (WP_000343449.1) MscS channels. All the analysis was performed in Clustal Omega (https://www.ebi.ac.uk/Tools/msa/clustalo/). Linear topology predictions were done comparing the transmembrane domain predictions of TopPred, TMpred and TMHMM 2.0 (https://www.expasy.org/tools/). Sequence similarity was analyzed using SIAS online tool (http://imed.med.ucm.es/Tools/sias.html). BLOSUM62, PAM250 and GONNET indexes were used to compare TcMscS and other MscS-like protein sequences.

*Molecular simulation of TcMscS structure*

Structural predictions were performed by six independent servers/predictors (I-TASSER, Robetta, Phyre2, SWISS-MODEL, IntFOLDTS, and RaptorX). Eight different structures of *E. coli* MscS (2oau, 2vv5, 4agf, 4hwa, 5aji) and its homologs from *T. tengcongensis* (3t9n TtMscS, 3udc chimera of TtMscS and EcMscS beta barrel), and from *H. pylori* (4hw9) were selected as templates based on automated analysis of sequence similarity and structural trends. All the monomeric predictions were assembled into heptamers using SymmDoc. After visual and automated inspection of the models, comparison of the consensus trends, and filtering for matching the hydrophobicity pattern of the membrane, and absence of severe structural conflicts we have picked one of the models (produced by IntFOLD2 server) as the most likely draft for the closed state conformation of TcMscS. We have manually adjusted the C-terminal amphipathic helix (connected by long flexible linkers to the cytoplasmic beta domains) to form a coiled-coil barrel, similar to what was observed in the crystal structures of another type of bacterial mechanosensitive channel, MscL.

The whole assembly was embedded into pre-equilibrated POPC bilayer and refined in Molecular Dynamics simulations (CHARMM36 force field) using iterative relaxation (5 ns) / symmetry-driven simulated annealing (1 ns) protocol. The refined closed state model was taken as a starting point in search of the open state structure using MD simulations with Extrapolated Motion Protocol to explore the conformational space. Structural analysis of the produced 5400 structures revealed the most promising candidate for the open state, included in Fig. 1.

*TcMscS cloning, expression and antibody generation*

The complete ORF of *TcMscS* was amplified with primers 1 and 2 (Table S1) from total genomic DNA from Y strain epimastigotes, cloned into pCR$^{TM}$-Blunt II TOPO® vector (Invitrogen), and sub-cloned into the pQE80 bacterial expression vector with restriction sites BamHI and HindIII. Bacterial expression of *TcMscS* ORF in BL21 plysS was induced overnight at 37°C with 1 mM isopropyl β-D-1-thiogalactopyranoside (IPTG). Recombinantly expressed TcMscS protein was extracted under denaturing conditions and purified with a nickel-agarose affinity column (Pierce). Polyclonal antibodies against the whole protein were obtained by guinea pig immunization, following standard protocols (Cocalico Biologicals Inc., Reamstown, PA).

*Western-blot analysis*

For western blot analysis, parasites were collected at 1,600 x *g* for 10 min, washed twice in PBS, pH 7.4, and resuspended in modified RIPA buffer (150 mM NaCl, 20 mM Tris-Cl pH 7.5, 1 mM EDTA, 1% SDS and 0.1% Triton X-100) containing protease inhibitor cocktail. Total homogenates were separated by SDS-PAGE, transferred onto nitrocellulose membranes and blocked overnight with 5% nonfat dry milk in PBS-0.1% Tween 20 (PBS-T). Blotting was done with anti-TcMscS (1:1,000) or anti-myc (1:250). Secondary antibodies conjugated with horseradish peroxidase or fluorescent DyLight 680 or 800 (LI-COR, Inc., Lincoln, NE) were used when indicated. Membranes were stripped with 62.5 mM Tris-HCl, pH 6.8, 2% SDS, 1% βmercaptoethanol at 50°C for 30 min, washed in PBS-T and incubated with monoclonal α-tubulin (Sigma) (1:5,000) as a loading control.

*Immunofluorescence analysis*

Epimastigotes, bloodstream trypomastigotes and host cells containing intracellular amastigotes were fixed for 30 minutes in 4% paraformaldehyde. Fixed cells were attached to poly-L-lysine-treated glass coverslips for 10 minutes. Samples were permeabilized with 0.3 % Triton X-100 for 3 minutes, washed in 1x PBS three times and incubated in 50 mM NH4Cl for 30 minutes at room temperature. After blocking overnight at 4 °C in 3 % bovine serum albumin (BSA) solution, the cells were incubated with antibodies against TcMscS (1:500), , actin (1;250), tubulin (1:500), SSP1 (1:250) or SSP4 (1:250) as indicated. Anti-SSP1 (# NR-50891) and anti-SSP4 (# NR050892) were obtained from BEI Resources, NIAID, NIH. The secondary antibodies were conjugated with Alexa- fluor 488 or 594 (1:3,000) (Thermo Fischer Scientific, Inc., Waltham, MA). Coverslips were mounted with Fluoromount-G® (SouthernBiotech, Birmingham, AL) containing DAPI (5 μg/mL). To evaluate the localization of TcMscS under osmotic stress, epimastigotes exposed to hypo or hyper osmotic conditions were fixed at times 0, 2 and 5 minutes post-stress and processed for immunofluorescence analysis as indicated before. Immunofluorescence samples were imaged in an Olympus© IX83 inverted microscope system and processed with CellSense Olympus software. Alternatively, samples were imaged in an ELYRA S1 (SR-SIM) Super Resolution Microscope (Zeiss) and processed with ZEN 2011 software, kindly provided by the Biomedical Microscopy Core at the University of Georgia.

### 2) Supplemental Results

**Video 1. Motility of Cas9 epimastigotes.**

**Video 2. Motility of TcMscS-KO epimastigotes.**

***Supplemental Tables.***

**Table S1. Sequences of primers used.**

**Table S2. Changes in cell volume upon hypoosmotic stress.**

**Table S3. Changes in cell volume upon hyperosmotic stress.**

**Table S4. Quantification of intracellular amastigotes.**

**Table S5. Quantification of extracellular trypomastigotes.**

***Supplemental Figures.***

**Figure S1. Sequence alignment of TcMscS.** Full protein sequence alignment of four MscS-type channels from *T. cruzi T. cruzi* Esmeraldo-like (TcCLB.504171.40) and non-Esmeraldo-like (TcCLB.509795.40) haplotypes of the CL Brener reference strain, *T. brucei* (Tb427.10.9030) and *L. major* (LmjF.36.5770), with two bacterial MscS channels from *E. coli* (WP_000389819) and *H. pilory* (WP_000343449.1). The position of the transmembrane domains TM1, TM2 and TM3 are underlined, the position of the putative gate residues is indicated by red arrows, conserved residues forming the hinges of TM3 are indicated with asterisks.

**Figure S2. TcMscS targeting by CRISPR/Cas9 produced a knockdown effect. A.** Partial alignment of TcMscS nucleotide sequencing (top panel) and predicted protein (bottom panel) obtained from wild-type epimastigotes CL Brener and Y strains, parasites expressing a scrambled sgRNA and Cas9 and those in which TcMscS specific sgRNA2 was transfected (TcMcsS-KD). A single nucleotide deletion in position 178 results in a shift in the ORF (black arrow) and a premature stop codon (red arrow). **B.** Growth curve of epimastigotes WT (black line), Cas9 (red line) and TcMscS-KD (blue line). Mean ± SEM of 3 independent experiments in triplicate (** p<0.01). **C.** RT-qPCR quantifying the expression of TcMscS in epimastigotes control (Cas9) vs TcMscS-KD. The values are indicated as $\Delta\Delta$Cq respect to the expression in WT epimastigotes and normalized against GAPDH as housekeeping gene. All the samples were collected from parasites at 4 days of growth. The values are Mean $\pm$ SEM of 3 independent experiments in triplicate (# *p = 0.040*). **D.** Immunofluorescence analysis of TcMscS (red) in epimastigotes Y strain (WT), Cas9 and TcMscS-KD. The expression of Cas9 was detected with anti-GFP antibodies. Nuclei and kinetoplasts were DAPI stained. The position of the contractile vacuole is indicated with yellow arrows. Bar size= 5 µm.

**Figure S3. TcMscS expression levels in epimastigotes and trypomastigotes. A.** Western blot analysis of epimastigotes with anti-TcMscS antibodies. Whole cell lysates were obtained from Cas9 expressing parasites, wild type Y strain (WT), TcMscS-KD (KD) and TcMscS-KO (KO). The purified recombinant protein was used as a positive control. Tubulin was used as loading control for all the samples. **B.** Quantification of TcMscS expression levels in epimastigotes by RT-qPCR. All the samples were collected from parasites at 4 days of growth. **C.** TcMscS expression levels in trypomastigotes by RT-qPCR. The values are indicated as $\Delta\Delta$Cq respect to the expression in WT and normalized against GAPDH as housekeeping gene. Mean $\pm$ SEM of 3 independent experiments in triplicate (** p = 0.040, ** p= 0.0001, # p= 6.43E-07*).

**Figure S4. Complementation of TcMscS-KO strains. A. Western blot analysis.** Total homogenates of WT, KO, complemented strains C1 and C2 carrying TcMscS-PAM-mutated-myc and C3 expressing TbMscS-myc were analyzed with anti-myc antibodies. **B. Growth curve** of epimastigotes WT (solid black line), KO (green line) and complemented strains C1 (orange line), C2 (dashed black line) and C3 (purple line). The values are Mean $\pm$ SD of 3 independent experiments in triplicate. **C.** Immunofluorescence analysis of complemented strains. Morphology and localization of myc-tagged TcMcsS (C1 and C2) and TbMscS (C3) where analyzed with anti-myc antibodies (green). TcMscS-KO cells were used as controls. Nuclei and kinetoplasts were DAPI stained. Bar size: 2 µm.

**Figure S5. Localization of TcMscS in epimastigotes under osmotic stress conditions.** Representative images showing the localization of TcMscS in epimastigotes under isosmotic, hypoosmotic or hyperosmotic conditions and different time points after stress. TcMscS was detected with specific antibodies against the channel. Nuclei and kinetoplasts were DAPI stained. Bar size: 5 µm.

**Figure S6. Volume of TcMscS mutants under isosmotic conditions.** Epimastigotes cell volume under isosmotic conditions was calculated based in the absorbance at 550 nm. The values were normalized respect to the values of wild type parasites (WT) and expressed as relative units. Values are expressed as mean ± SEM of 7 independent experiments (* $p < 0.001$).
